# Germline-targeting HIV immunogen induces cross-neutralizing antibodies in outbred macaques

**DOI:** 10.1101/2025.10.22.684023

**Authors:** Nitesh Mishra, Bo Liang, Ryan S. Roark, Amrit Ghosh, Sean Callaghan, Wen-Hsin Lee, Xuduo Li, Anh L. Vo, Gabriel Avillion, Rohan Roy Chowdhury, Rumi H. Habib, Frederic Bibollet-Ruche, Gabriella Giese, Prabhgun Oberoi, Khaled Amereh, Anjali Somanathan, Yuxin Zhu, Yuexiu Zhang, Muzaffer Kassab, Lifei Tjio, Sharaf Andrabi, Raphael A. Reyes, Joel D. Allen, Nicole E. James, Kipchoge N. Randall, Lara van der Maas, Elana Ben-Akiva, Kasia Kacmarek-Michaels, Samantha Plante, Christian L. Martella, Ashwin N. Skelly, Ajay Singh, Jonathan Hurtado, Katharina Dueker, Tazio Capozzola, Rebecca Nedellec, Gabriel Ozorowski, Mark M. Lewis, Samantha Falcone, Andrea Carfi, Sunny Himansu, Lawrence Shapiro, Max Crispin, Beatrice H. Hahn, Bryan Briney, Darrell J. Irvine, Dennis R. Burton, Andrew B. Ward, Facundo D. Batista, Peter D. Kwong, George M. Shaw, Andrabi Raiees

## Abstract

Germline-targeting-(GT) is a promising strategy to activate rare broadly neutralizing antibody (bnAb)-producing B cells against HIV, but induction of such responses in outbred animals has not been achieved. Using antibody-guided structure-based design, we engineered a germline-targeting trimer immunogen Q23-APEX-GT2 that primes diverse V2-apex bnAb precursors. Q23-APEX-GT2 efficiently activated V2-apex-specific B cells in humanized knock-in mice and consistently elicited immunofocused antibody responses in rhesus macaques, priming multiple long CDRH3-loop bnAb-B cell lineages. Monoclonal antibodies from immunized macaques exhibited broad heterologous HIV trimer binding and cross-neutralization. Atomic-level structural studies confirmed precise epitope targeting and revealed CDRH3-paratope configurations that mirrored those of human V2-apex bnAbs. This study provides proof-of-principle for successful priming and maturation of authentic V2-apex bnAb precursors in outbred macaques, underscoring the potential of V2-apex-targeted vaccines.

**HIGHLIGHTS:**

- Engineered Q23-APEX-GT2 trimer to stimulate diverse V2-apex bnAb B cell precursors
- Q23-APEX-GT2 primed rare V2-apex bnAb B cells in mice and outbred rhesus macaques
- Q23-APEX-GT2 elicited immunofocused antibody responses and diverse V2-apex B cell lineages with desirable long-CDRH3 paratope properties
- Q23-APEX-GT2 alone induced V2-apex antibodies with broad HIV trimer binding and modest neutralization breadth
- Structural analysis confirmed bnAb site targeting, mirroring human and rhesus V2-apex bnAbs

## INTRODUCTION

A major goal of HIV vaccine research is to elicit broadly neutralizing antibodies (bnAbs), which have been shown to be protective both in non-human primate and human studies ^1–4^. Although some HIV-infected individuals eventually develop bnAbs, this occurs infrequently ^5–11^ due, at least in part, to bnAbs being encoded by rare B cell precursors and requiring complex affinity maturation pathways ^7,12–19^, making their induction through vaccination especially challenging. Accordingly, many recent HIV vaccine strategies have focused on activating rare bnAb-producing B cell precursors via germline-targeting (GT) immunogens, followed by rationally designed boosting strategies ^18,20–29^. This approach has demonstrated promise in priming bnAb B cell precursors in both preclinical animal models and human trials ^30–39^, although no studies to date have induced bnAbs, either in B memory cells or secreted antibody, in stringent outbred animal models.

One of the most promising targets for HIV vaccines is the V2-apex bnAb site ^13,27,28,40–45^. Antibodies targeting this epitope are typically potent and broad, occur early in infection, require relatively low levels of somatic mutations, and are among the most common to occur in natural HIV infection ^5,6,8,45–52^. These V2-apex bnAbs target a lysine-rich patch on the V2 C-strand and neighboring glycans near the three-fold axis of the Env trimer, making these antibodies generally trimer-specific ^28,41,53–57^. A critical feature of these bnAbs is a long anionic heavy-chain complementarity-determining region 3 (CDRH3), which enables them to penetrate the glycan shield and reach the underlying cationic protein surface ^41,55,56,58^. However, a significant barrier to eliciting V2-apex bnAbs is the rarity of human B cell precursors with long CDRH3 loops ^13,59,60^. Nonetheless, these rare bnAb precursors possess germline D-gene-encoded paratope features that could be exploited by targeted vaccines, making a germline-targeting approach potentially very effective ^13,28,41,43^. We hypothesized that the key obstacle to the elicitation of V2-apex bnAbs is the development of trimer immunogens that closely mirror the native Env of wildtype infectious virions but also contain select engineered mutations that substantially enhance their affinity for diverse germline B cell bnAb precursors, or unmutated common ancestors (UCAs). This strategy ensures that germline B cell precursors that have the potential for development into actual bnAbs are preferentially primed, and once activated, affinity-mature to bind native-like Env structures including those on heterologous viruses.

In this study, by using antibody-guided structure-based reverse vaccine design, we generated a germline-targeting trimer immunogen (Q23.APEX-GT2) to activate rare V2-apex bnAb B cell precursors. Immunization in outbred rhesus macaques elicited a consistent antibody response highly focused to the HIV Env V2-apex bnAb site. Isolation of monoclonal antibodies (mAbs) from memory B cells showed the expansion of diverse V2-apex bnAb site-targeting long CDRH3 B cells across multiple vaccinated macaques. These mAbs bound a variety of heterologous Env trimers representing the global HIV diversity. A subset of these mAbs exhibited modest neutralization breadth against tier 2 heterologous viruses, which to our knowledge is the first example of consistent heterologous neutralization induced by vaccination in outbred macaques. Atomic-level structural studies of the isolated antibodies confirmed targeting of the V2-apex bnAb site in diverse binding modes mirroring those observed in human and rhesus V2-apex bnAbs elicited by HIV and SHIV infection ^41,53,55–57,61^. Thus, our study is the first to demonstrate the induction authentic V2-apex bnAb precursors and their partial maturation into cross-neutralizing antibodies in an outbred monkey model using a germline-targeting immunogen approach. Notably, this occurred after a homologous prime and boost vaccination with a single germline-targeting trimer immunogen. Overall, the results suggest that V2-apex bnAb elicitation may be relatively straightforward, requiring limited boosting and few somatic hypermutations, if the appropriate rare long CDRH3 loop B cells can be effectively primed.

## RESULTS

### Rational design of Q23-APEX-GT2 with enhanced binding to V2-apex bnAb UCAs

Germline-targeting (GT) has emerged as a promising approach in HIV-1 vaccine design, aiming to activate rare broadly neutralizing antibody (bnAb) precursors by engineering high-affinity immunogens. However, many GT strategies focus narrowly on engaging only certain antibody lineages, which may limit the breadth of response against diverse virus strains. In contrast, we employed a modified GT-approach designed to target a wider array of HIV Env V2-apex bnAb precursors, irrespective of their paratope structural diversity. To this end, we previously showed that V2-apex bnAbs possess germline D-gene-encoded anionic residues (YYD) that enable them to interact readily with the positively charged V2-apex core bnAb epitope ^28,41^. We also demonstrated that a unique set of HIV Envs naturally have a rare glycan hole that facilitates binding of V2-apex bnAb unmutated common ancestors (UCAs) in the native-like trimer configurations ^43^. When these unique Envs were delivered into rhesus macaques as SHIVs in an infection model, one of them, Q23.17, reproducibly induced V2-apex bnAbs, indicating an inherent propensity to elicit V2-apex site-targeted bnAb responses ^61,62^.

Using an antibody-guided structure-based design approach, we recently developed a Q23.17-based, prefusion-stabilized, V2-apex germline-targeted trimer (Q23-APEX-GT1) that exhibited native-like trimer properties and primed rhesus V2-apex bnAb UCA knock-in B cells, resulting in moderate neutralization breadth and potency (Ghosh et al., 2025 *in preparation*). Here, we utilized a large panel of *bona fide* V2-apex bnAb UCAs from SHIV-infected rhesus macaques^62^, along with molecular information on how V2-apex bnAbs bind to HIV trimers (Figure S1) ^61^, to improve the broad UCA binding properties of Q23-APEX-GT1. Our strategy was to prime a wider array of V2-apex bnAb precursor antibodies, potentially expanding the precursor pool for the development of bnAbs against HIV.

Starting with the recently stabilized Q23-APEX-GT1 trimer as a base construct, we generated a large number of variants by introducing additional antibody-trimer structure-based amino acid mutations alone and in combination within the V1V2 region at residues 130, 132, 135, 158, 167, 169, 170, 171, 173 (HXB2 reference numbering) (Figure 1A-B, S1). By testing the binding of these mutants to diverse V2-apex bnAb UCAs, we identified Q23-APEX-GT1-T132R-S158T (designated Q23-APEX-GT2) as the most improved construct, since it bound 8 of 10 rhesus and human V2-apex bnAb UCAs with high affinity (Figure 1B-C). The optimized Q23-APEX-GT2 trimer showed substantial KD affinity improvements (Figure 1C, S2), driven primarily by enhanced on-rate constants (Kon) (p < 0.05), highlighting its improved germline precursor engagement—a key feature for germline-targeting immunogens. Since the S158T mutation changes the N156 glycan sequon from NxS to NxT, we performed site-specific glycan analysis of Q23-APEX-GT2 using mass spectrometry to determine the glycan composition and occupancy. Overall, the glycan profile of Q23-APEX-GT2 was similar to that of the base construct Q23-APEX-GT1, except for heterogeneity in glycan maturation (predominantly displaying mannose-rich glycans) and slightly reduced occupancy (up to 7%) at glycan positions N156, N160 and N197 (Figure 1D-E, S3). The abundance of oligomannose-type glycans further indicates the assembly of a well folded native-like trimer ^63^. The antigenicity of Q23-APEX-GT2 was similar to that of Q23-APEX-GT1 as demonstrated by the binding profiles of a panel of HIV bnAbs and non-neutralizing antibodies (nnAbs) targeting various Env epitopes (Figure S3A). This conservation of glycan presentation and native-like trimer antigenicity supports the structural integrity of the engineered Q23-APEX-GT2 trimer. Finally, we tested a membrane-bound version of Q23-APEX-GT2 as a potential mRNA-based immunogen and found that it exhibited a similar binding profile to V2-apex precursors and mature bnAbs compared to the soluble protein, with no negative impact on its overall antigenic profile (Figure S2C, S3). Q23-APEX-GT2 maintained an HIV tier 2 Env antigenic and neutralization sensitivity profile, demonstrating binding and neutralization by bnAbs targeting canonical Env bnAb sites while showing minimal to no binding or neutralization by non-neutralizing antibodies (nnAbs) that recognize “open” Env structures (Figure S3).

**Figure 1.**
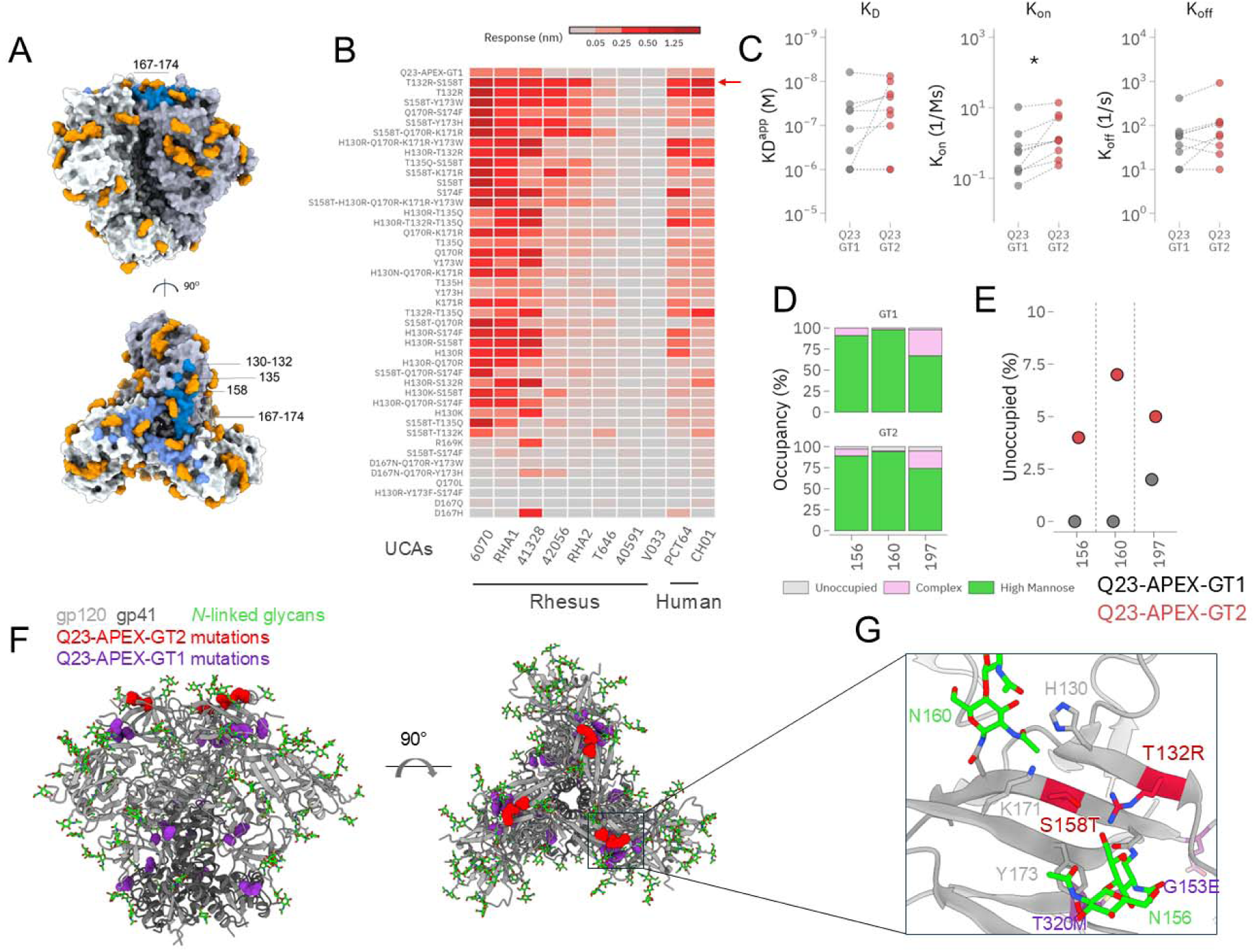
Antibody-guided structure-based design of Q23-APEX-GT2 for enhanced binding to diverse prototype V2-apex bnAb precursors. See also Figures S1, S2, S3 and S4. A. V1V2 residues (130, 132, 158, 167-174) that potentially interact with V2-apex bnAbs, and/or their precursors are shown on crystal structure of Q23 DS-SOSIP (PDB: 7LLK). Residues are colored blue while glycans are shown in orange. B. Heat map representing the BLI binding response of a broad V2-apex UCA panel against V1V2 mutants generated on Q23-SCT27 or Q23-APEX-GT1 backbone. First row in the heat map shows the binding against the base construct (Q23-APEX-GT1 or GT1). Mutants are ordered based on average binding across the UCA panel. 9 rhesus (6970, RHA1, 41328, 42056, P1O21, RHA2, T646, 40591 and V033) and 2 human (PCT64 and CH01) UCAs were included in the BLI binding screen. Based on broad binding to UCAs, Q23-APEX-GT1-T132R-S158T variant, designated as Q23-APEX-GT2 (or GT2 – indicated by red arrow) was down-selected. C. Apparent binding affinity constants (K_D_), on-rate constants (K_on_) and off-rate constants (K_off_) of base construct (GT1) compared to germline-targeting lead candidate (GT2). A significant increase in K_on_ was observed (p < 0.05). D. Glycan occupancy assessed by proteomics-based site-specific glycan analysis (SSGA) at key V2-apex glycans (position 156, 160 and 197) in GT1 and GT2. High mannose in faded green; complex glycan is shown in light pink; unoccupied fraction is shown in gray. E. Increased unoccupied fractions at position 156, 160 and 197 seen for GT2. F. Cryo-EM structure of germline-targeting immunogen Q23-APEX-GT2 showing a compact assembly and trimeric apex. gp120 is colored light gray, gp41 is colored dark gray, glycans are colored green. GT1 mutations are shown in violet. GT2 mutations are at the apex and shown in red. G. Close-up view of potential interactions between key GT2 mutations 132R with the N156 glycans at the apex of GT2.

To confirm a native-like trimer conformation, we resolved the structure of Q23-APEX-GT2 at 2.9 Å resolution using single-particle cryogenic electron microscopy (cryo-EM). This analysis revealed a compact trimeric apex consistent with the prefusion-closed conformation of Env (Figure 1F, S4). Further examination of the local environment of engineered residues T132R and S158T revealed the introduction of an interactive network spanning residues 132, 156, 158, 171, and the first *N*-acetylglucosamine residue of N156-glycan, suggesting that fortification of interactions across the A-, B-, and C-strands of the trimer apex contributes to the observed enhanced antibody binding (Figure 1G and S4). Trimer apex-stabilization was also achieved through the initial Q23-APEX-GT1 modifications, including the T320M mutation which fills a hydrophobic pocket lined by V2 residues 154, 175, and 177, and the G153E mutation, which is known to suppress exposure of the V3 loop (Figure S4). Thus, the structural data suggest a synergy between the apex-stabilizing mutations of GT1 and the breadth-enhancing mutations of GT2, which together are likely responsible for the improved binding of the Q23-APEX-GT2 trimer to a broad spectrum of human and rhesus V2-apex bnAb precursors, regardless of their structural class ^61^.

### The engineered Q23-APEX-GT2 trimer efficiently primes rare human V2-apex bnAb UCA knock-in B cell precursors

To evaluate the *in vivo* priming efficacy of engineered Q23-APEX-GT2, we tested its ability to engage and expand precursors of a defined V2-apex bnAb lineage using a knock-in mouse model. We selected a mouse model with B cells bearing the least mutated common ancestor (LMCA) heavy chain (IGH) and light chain (IGK) of bnAb PCT64, which was isolated from a human donor, as the PCT64 class is among the most prevalent in the human naïve B cell repertoire ^13,42,46^. Given the high frequency of PCT64^LMCA^ B cells in our KI mouse model, we used an adoptive transfer system where (1×10^5^) CD45.2^+^ PCT64^LMCA^ B cells were transferred into congenic CD45.1^+^ C57BL/6J recipient mice to decrease precursor frequency to try to mimic the rare precursor frequency observed in humans. This adoptive transfer resulted in approximately 20 antigen-specific CD45.2 B cells per 1×10^6^ B cells in spleen^42^. Cohorts of these adoptively transferred mice were immunized with: SMNP adjuvant alone (Arm 1 – control), Q23-APEX-GT2 trimer protein + SMNP (Arm 2) and membrane-anchored Q23-APEX-GT2 trimer mRNA lipid nanoparticles (Arm 3) (Figure 2A). To assess the priming efficiency of Q23-APEX-GT2, we harvested draining lymph nodes (LNs) at weeks 2 and 4 post-vaccination and analyzed antigen-specific B cell responses via flow cytometry (Figure 2B-C). Enhanced LN germinal center (GC: CD95^hi^CD38^lo^) B cell responses were observed in both Q23-APEX-GT2 protein and mRNA groups compared to the adjuvant-only control. At week 2, the protein group showed a significantly higher GC B cell response compared with the mRNA group (20% vs 8%), but by week 4 both groups were comparable (7% vs 9%). A substantial proportion of LN GC B cells were PCT64^LMCA^ CD45.2 B cells (20% vs 2%) at both time points, indicating efficient priming, expansion and retention of the rare human V2-apex bnAb precursors in the GCs (Figure 2B-C). Consistent with GC formation, we observed that immune sera collected at weeks 2 and 4 post-prime exhibited Q23-APEX-GT2-specific antibody binding (Figure 2D). A substantial fraction targeted the V2-apex bnAb site, as confirmed by ELISA binding to Q23-APEX-GT2 trimer and its N160K variant lacking the critical N160 glycan; N160K substitution abrogates PCT64 and other V2-apex antibody binding ^28,42,46^.

**Figure 2.**
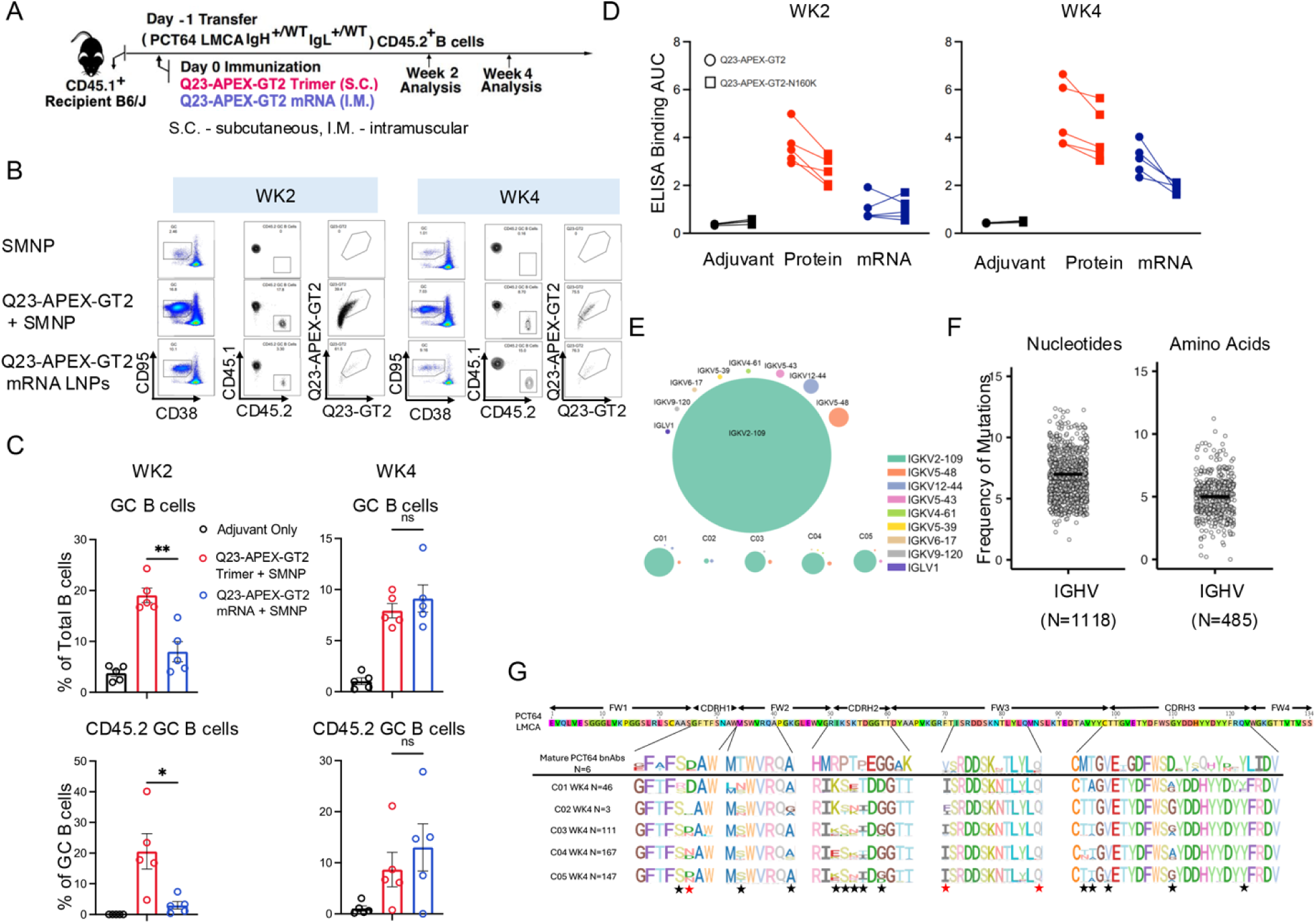
Q23-APEX-GT2 trimer successfully primes human V2-apex bnAb precursor PCT64^LMCA^ encoding rare B cells. A. Schematic representation of PCT64^LMCA^ mouse immunization studies. 1 × 10^5^ B cells of CD45.2-encoded PCT64^LMCA^ precursors were injected intravenously into the tail vein of CD45.1 WT mice recipients (B6.SJL-Ptprca Pepcb/BoyJ). Post adoptive transfer mice were immunized with SMNP adjuvant [(5μg) (Arm 1 - control)], Q23-APEX-GT2 protein + adjuvant [(20μg protein + 5μg adjuvant) (Arm 2)] and membrane-anchored Q23-APEX-GT2 mRNA lipid nanoparticle (LNPs) [(1μg mRNA LNPs) (Arm 3)]. The adjuvant or protein + adjuvant was administered subcutaneously and the mRNA LNPs were injected by intramuscular route. Lymph nodes and immune sera were collected at week 2 and 4 time points. B. Representative flow cytometry plots of B cells from draining LN at 2- and 4-weeks post immunization. Germinal Center (GC) B cell and CD45.2 GC B cell responses to Q23-APEX-GT2 trimer in weeks 2 (left) and 4 (right) post immunization are shown. C. Quantification of GC B cells and CD45.2 GC B cells in week 2 (left) and week 4 (right) post immunization with SMNP adjuvant only, Q23-APEX-GT2 trimer protein with SMNP and Q23-APEX-GT2 mRNA LNPs (n=5). Data is represented as mean ± SEM. D. The ELISA binding AUC titers of serum binding to Q23-APEX-GT2 and the V2-apex epitope knock-out (N160K) at wk2 (left) and WK4 (right) post-immunization. E. Distribution of IG Kappa/Lambda V gene usage in Q23-APEX-GT2 protein vaccinated individual mice (C01–C05). The antibody sequences were derived from LN B cells harvested at week 4 post immunization. Data are shown as bubble plots, where the circle size represents its proportion of the total V gene count. Light chain enrichment breakdown by individual mice in sublots. F. Somatic hypermutation (SHM) levels in the Q23-APEX-GT2 vaccination derived PCT64^LMCA^ heavy chains. SHM levels were calculated using both nucleotide and amino acid sequences of the v-gene. Each dot represents a sequence compared to the PCT64 germline, with the black line indicating the mean number of mutations. G. Sequence logograms across individual mice obtained from week 4 (bottom 5 rows) of highly mutated residues, compared with the least mutated common ancestor (LMCA, top line) of PCT64 and six mature PCT64 bnAbs. SHMs unique to mouse immunization (black) and shared with PCT64 mature bnAbs (red). p values were calculated by a Mann-Whitney test (A and H). *p < 0.05; ns, not significant. Error bars are SEM

To assess whether PCT64^LMCA^ B cells underwent somatic hypermutation (SHM) and acquired PCT64-like mutations following Q23-APEX-GT2 immunization, we isolated class-switched Q23 APEX-GT2+ PCT64^LMCA^ B cells at week 4 post immunization for 10X B cell receptor (BCR) sequencing from five protein-vaccinated mice (C01–C05). Sequence analysis revealed a strong selection for the IGKV2-109 mouse light chain paired with the PCT64^LMCA^ heavy chain in all five mice (Figure 2E). Analysis of PCT64^LMCA^ IGH lineage evolution revealed extensive diversification by week 4 (Figure 2F, G). By week 4 post-immunization, substantial SHM were observed in the IGH V region (∼5% amino acid level). Mutations accumulated in recurrent positions within HCDR1, HCDR2, and HCDR3 over time. Further analysis showed antigen-driven convergent SHM patterns across mice, with some mutations occurring at positions shared with mature PCT64 bnAbs (Figure 2G). Overall, Q23-APEX-GT2 effectively primed and expanded rare human PCT64^LMCA^ B cell precursors, facilitating germinal center formation, inducing V2-apex-targeted antibody titers, and promoting B cell maturation along favorable evolutionary pathways.

### Q23-APEX-GT2 trimer elicits V2-apex immunofocused antibody responses in outbred rhesus macaques

The rhesus macaque model provides a more competitive B cell environment in an outbred setting, making it a more physiologically relevant system to evaluate the ability of Q23-APEX-GT2 to prime and mature rare V2-apex-targeting B cell precursors. To assess the immunogenicity of the Q23-APEX-GT2 trimer in RMs, two groups (6 animals per group) were subcutaneously immunized with either 100 µg of trimer protein plus saponin/MPLA nanoparticle adjuvant (SMNP) ^64^ using a slow-delivery escalating-dose protocol (Arm-1) ^65,66^, or 100 µg of trimer mRNA lipid nanoparticles (LNPs) via bolus injection (Arm-2) (Figure 3A). Both groups also received a bolus boost of 100 ug of the same immunogen at week 10. Plasma samples were collected at various time points, and excisional lymph node biopsies were performed at weeks, −6, 4, 8, 12, and 14 to monitor antibody and B cell responses.

**Figure 3.**
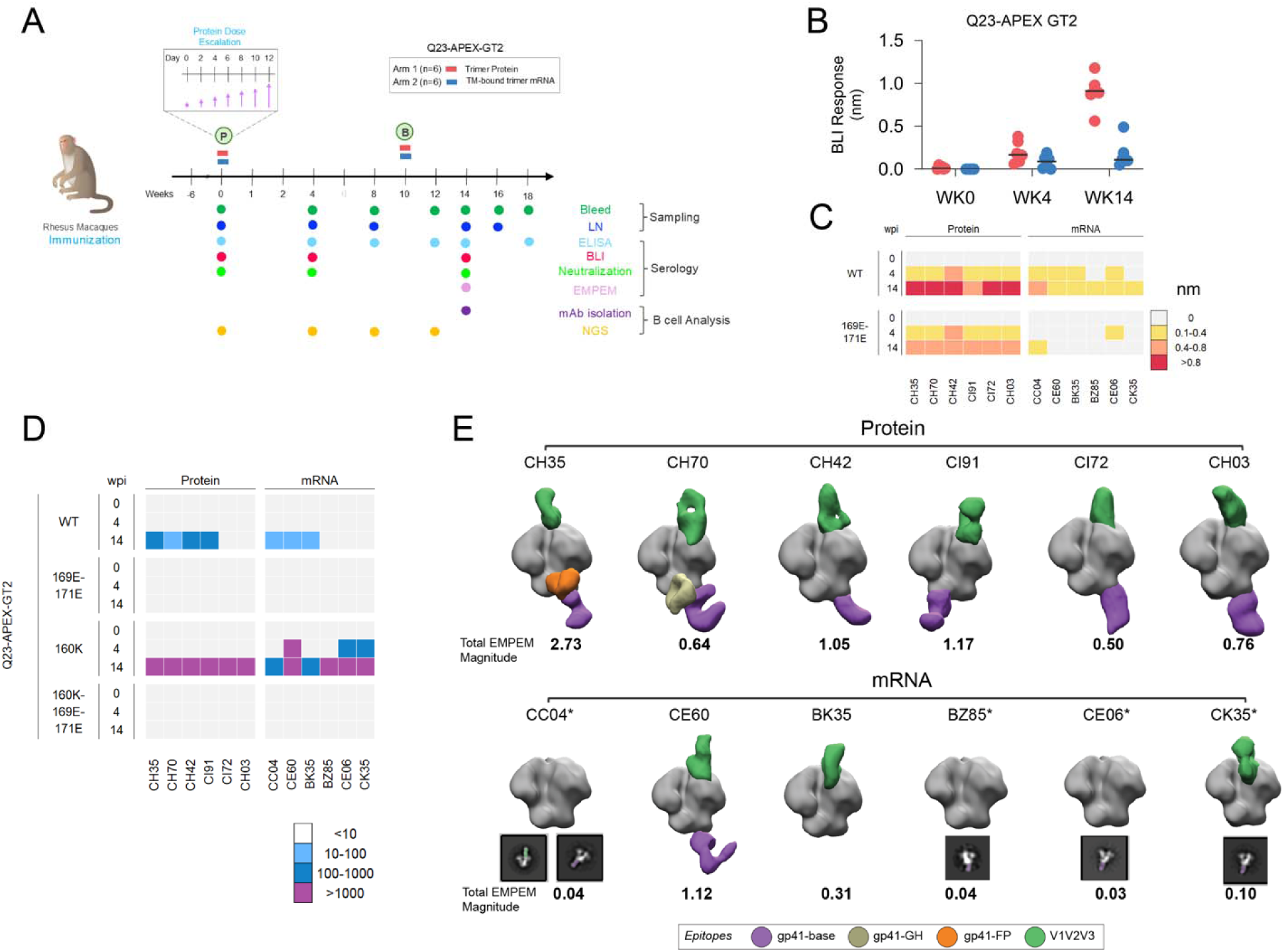
Q23-APEX-GT2 trimer immunization in rhesus macaques and analysis of serum antibody responses. See also Figures S5, S6 and S7. A. Schematic showing rhesus macaque immunization with Q23-APEX-GT2 trimer protein + SMNP (Arm 1) and membrane-anchored mRNA LNPs (Arm 2). Rhesus macaques were primed at week 0 with 100 μg Q23-APEX-GT2 protein + SMNP adjuvant (Arm 1) or 100 μg Q23-APEX-GT2 mRNA LNPs (Arm 2) administered subcutaneously and distributed into four injection sites (25 μg each). The Arm 1 protein priming immunization was given as escalating dose (DE) over 2 weeks and a single bolus dose for mRNA arm. Both groups were bolus boosted with 100 μg of protein + SMNP (Arm 1) or mRNA LNPs (Arm 2). Serum, LN, and PBMC sampling longitudinally, as well as functional analyses, indicated by colored circles. B. Binding of polyclonal serum IgGs to Q23-APEX-GT2 trimer antigen by BioLayer Interferometry (BLI). Protein group is in red, mRNA group is in blue. The antigen specific antibody titers were detectable at week 4 post-prime in both groups and were boosted after week 1o boost. Stronger antibody titers were detected in the protein compared to the mRNA group. C. Heatmap showing binding of longitudinal polyclonal serum IgGs (weeks, 0, 4 and 14) to Q23-APEX-GT2 and its V2-apex epitope knockout (R169E-K171E) showing binding dependence on strand C. D. Heatmap showing serum neutralization of pseudovirus bearing the Q23-APEX-GT2 mutations (132R-158T) and several V2-apex bnAb epitope knockout virus variants. Neutralization is dependent on strand C residues as evident by loss of neutralization with 169E-171E variant. Increased neutralization by immune sera was observed upon160K glycan knockout and this was completely abrogated on strand C epitope knockout in the N160K backbone (160K-169E-171E), further highlighting strand C-dependent neutralization. E. Electron microscopy polyclonal epitope mapping (EMPEM) imaging of immune serum Fabs from Q23-APEX-GT2-immunized RMs highlights immunofocused response in all protein immunized animals and 3/6 mRNA immunized animals to Q23-APEXGT2 trimer. Composites of Three-dimensional (3D) reconstructions generated from the negative stain electron microscopy images of polyclonal serum antibody Fabs bound to Q23-APEXGT2 trimer are segmented. Fabs are colored based on targeted epitope region. Colored 2D classes are shown when too few particles were present for 3D reconstruction. Total EMPEM magnitude for each dataset is listed and described in more details in Methods.

Post-prime immune sera showed Q23-APEX-GT2-specific antibody responses, as measured by ELISA and BLI, with substantial increases after week 10 (Figure 3B, S5A-D). Higher antibody titers were observed in the adjuvanted protein group compared to the mRNA group (∼5-fold difference at WK14) (Figure 3B, S5A). Longitudinal serum antibody responses were further assessed for V2-apex epitope targeting by testing binding to strand-C residue substitutions (R169E-K171E (double knock-out - dKO)), which abolish binding of most V2-apex bnAbs ^28^, and binding to a glycan knockout at position 160 (N160K) on the Q23-APEX-GT2 trimer (Figure 3C, S5A-D). Positively charged V2-strand-C residues and the N160 glycan form the core binding region of the V2-apex bnAb site ^28^. Substitution of the V2-strand-C positively charged residues substantially reduced binding by immune sera (1.1 - 1.7-fold reduction for protein, complete abrogation for mRNA) in both protein- and mRNA-immunized groups, while responses against the N160 glycan KO variant were slightly enhanced (1.1 - 1.9-fold enhancement for mRNA) (Figure 3C, S5). Overall, the protein group induced stronger responses than mRNA, with evidence of targeting the strand C of the V2-apex bnAb site.

We also tested longitudinal serum neutralization against the immunogen-matched Q23-APEX-GT2 pseudovirus and its strand C dKO variant. 4 of 6 animals in the protein group and 3 of 6 in the mRNA group neutralized the autologous virus (40 - 430 ID50 range for protein, 10 - 75 ID50 for mRNA) in a V2-apex epitope-dependent manner (Figure 3D, S6). Consistent with binding data, mRNA-immunized animals had weaker nAb titers than the protein group (∼4.5-fold higher in protein group) (Figure 3D, S6). We also tested neutralization of immune sera using the non-germline-targeting wild-type (WT) Q23.17 tier-2 virus and its strand C- and N160 glycan-KO variants. Immune sera from one of the protein-group animals, CH35 showed weak but detectable neutralization of the WT Q23 virus (Figure S6). Notably, all animals in both groups developed high nAb titers (ID50 range: Protein, >21870 and mRNA, 260 - >21870) against the N160-deficient Q23.17 virus, with substantial increases following the week-10 boost, as detected in week-14 sera (Figure S6). These responses targeted the V2-apex strand C residues, as shown by the complete loss of neutralization against the N160-K169-K171 variant (Figure S6). No heterologous virus neutralization was observed (Figure S6). The results indicate that the Q23-APEX-GT2 trimer induces immunofocused polyclonal neutralizing antibody responses targeting the V2-apex bnAb site in outbred rhesus macaques.

To gain more insight into the epitopes targeted by polyclonal antibodies at week 14, we used electron microscopy polyclonal epitope mapping (EMPEM) ^67^. Antigen-binding fragments (Fabs) from the immune plasma of all 12 animals (6 from each group) were complexed with the Q23-APEX-GT2 trimer (Figure 3E, S7). Consistent with serum binding and neutralization data, EMPEM revealed that polyclonal antibody responses predominantly targeted the canonical V2-apex bnAb site in 6 of 6 protein-immunized animals and 3 of 6 mRNA-immunized animals (Figure 3E, S7). As expected, targeting of the trimer base was observed in the protein group but was largely absent in the mRNA group (Figure 3E, S7). The total EMPEM magnitude (average number of Fabs per trimer) was higher in the protein group compared to mRNA group (mean value of 1.14 for protein group and 0.27 for mRNA group; see also Methods and Figure 3E), which is consistent with the antibody titer results (Figure 3B and S5). These results support the above data to suggest that the Q23-APEX-GT2 trimer elicits V2-apex-focused antibody responses in outbred macaques.

### Q23-APEX-GT2 enriches long CDRH3 V2-apex bnAb site specific B cells

One objective of this study was to determine whether the Q23-APEX-GT2 trimer can prime V2-apex-targeting rare bnAb B cell precursors with long CDRH3 loops in outbred monkeys. To assess this, we used flow cytometry to isolate V2-apex-specific, IgG class-switched B cells from the lymph nodes (LNs) of six protein-vaccinated monkeys at week 14 or 16 (four or six weeks after the second immunization). To isolate epitope-specific B cells, we used Q23-APEX-GT2 trimer and dKO probes (Q23-APEX-GT2+/Q23-APEX-GT2-dKO-ve) as baits (Figure S8A), with antibody sequences obtained by sequencing the amplified heavy and light chains from antigen-sorted single B cells. Analysis of the week 14/16 lymph nodes showed that all 6 Q23-APEX-GT2 protein-vaccinated animals developed antigen-specific B cell responses (median = ∼3%), with a sizable fraction being V2-apex specific (median = ∼2.5%) (Figure S8B). We observed a strong enrichment of ≥22 amino acids CDRH3 loops (2- to 48-fold enrichment) in the V2-apex-specific B cells compared to the baseline naive B cell repertoire in RMs (Figure 4A), in all 6 vaccinated monkeys. These results demonstrate that Q23-APEX-GT2 immunization substantially shifted the distribution B cells with long CDRH3s (a signature feature for V2-apex bnAbs) in outbred RMs. The immunogenetic analysis of the isolated antibodies revealed that three major germline D genes, IGHD3-15, IGHD3-17, and IGHD3-18, contributed to the long CDRH3 mAbs, with IGHD3-15 being used in 5 of the 6 monkeys (Figure 4B). This preferential usage of the IGHD3-15 germline D gene in long CDRH3 V2-apex bnAbs is consistent with previous observations in rhesus bnAbs isolated from SHIV-infected monkeys ^57,61^.

**Figure 4.**
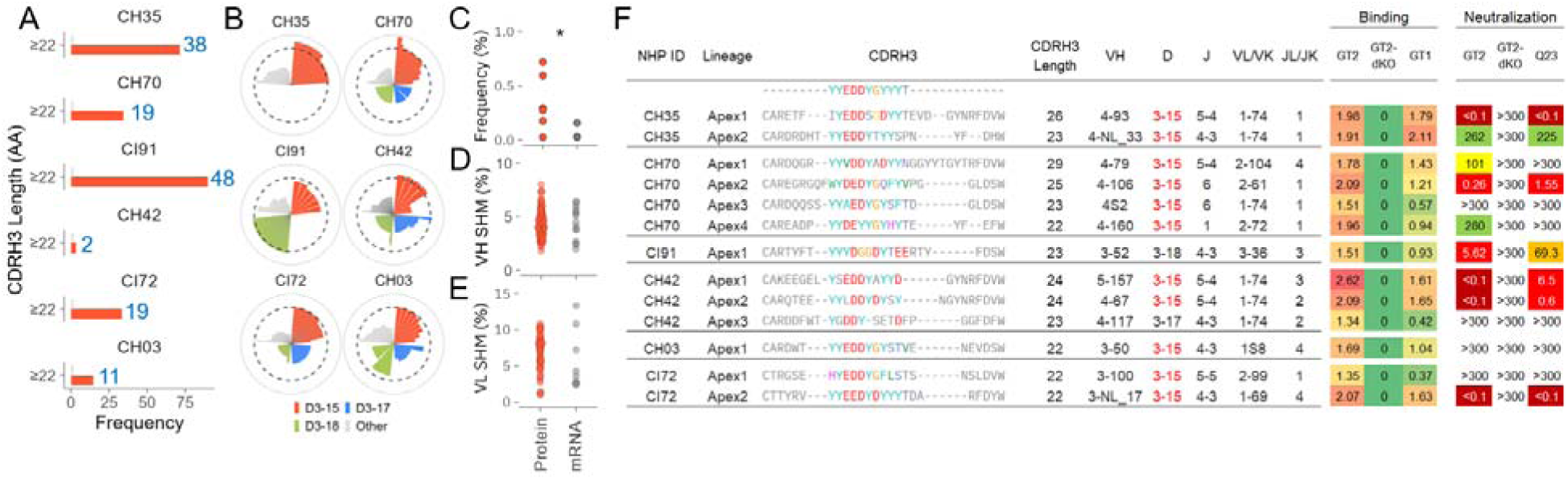
Isolation and functional characterization of Q23-APEX-GT2 trimer elicited rhesus V2-apex mAbs: immunogenetic features and epitope properties. See also Figure S8. A. Enrichment of Q23-APEX-GT2 V2-apex epitope sorted B cells from week 14 or 16 LNs of Q23-APEX-GT2 trimer protein immunized RMs. Bar plots show enrichment of long CDRH3s (length ≥22 AA) in protein-immunized animals in red; gray bar shows the frequency in naïve RMs (as a baseline reference). B. Circular bar plots showing the germline-D-gene usage in Q23-APEX-GT2 V2-apex epitope sorted B cells from panel A. IGHD3-15, IGHD3-17 and IGHD3-18 are colored while all other D-genes are combined as “others”. The dashed inner circle represents the CDRH3 length cutoff of ≥22 amino acids. C. Distribution of long CDRH3 B-cells (≥22AA) from Q23-APEX-GT2 V2-apex epitope sorted B cells in protein and mRNA immunized rhesus macaques. A significantly higher fractions of long CDRH3 B cells was observed in protein-immunized rhesus macaques. D. Somatic hypermutation in the heavy chain (HC) variable region of site-specific monoclonal antibodies (mAbs) from Q23-APEX-GT2 protein and mRNA immunized rhesus macaques, regardless of CDRH3 length. E. Somatic hypermutation levels in light chains (LC) of long CDRH3 (≥22 AA) mAbs from Q23-APEX-GT2 V2-apex epitope sorted B cells. F. CDRH3 sequence, length and gene assignments for V, D, J for heavy chain and V, J for light chains from representative long CDRH3 mAbs lineages from Q23-APEX-GT2 protein immunized rhesus macaques. Germline IGHD3-15 D-gene is shown in red and the germline encoded anionic CDRH3 residues are highlighted in colors. From each animal, representative sequences from each expanded lineage are shown. For each mAb, maximum BLI binding responses and IC_50_ neutralizations are shown with Q23-APEX-GT2 trimer protein and its corresponding virus and the strand C dKO variants and the WT Q23.17. All mAbs are dependent on V2-apex stand C core epitope.

**Figure 5.**
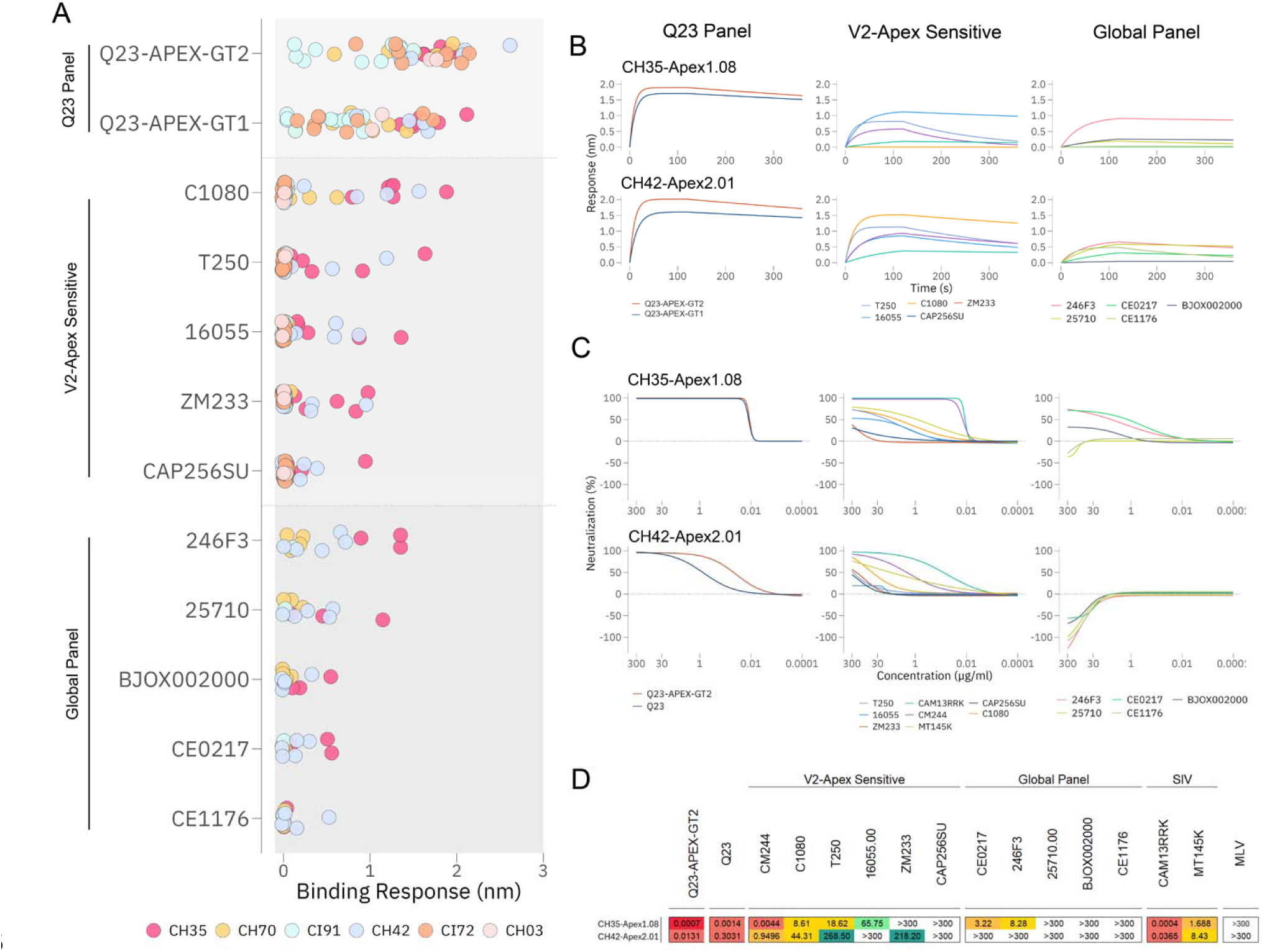
Q23-APEX-GT2 trimer elicited rhesus V2-apex mAbs exhibit cross-reactive binding with heterologous HIV trimers and modest neutralization breadth. See also Figure S8. A. Scatter plot illustrating BLI binding responses of all isolated mAbs to autologous (Q23-APEX-GT2 and Q23-APEX-GT1) HIV Envelope trimers, V2-apex neutralization-sensitive heterologous trimers (C1080, T250, 16055, ZM233, and CAP256.SU), and trimers from the global HIV-1 reference panel (246F3, 25710, BJOX002000, CE0217, and CE1176). Many Q23-APEX-GT2-elicited long CDRH3 rhesus V2-apex mAbs exhibited cross-reactive binding to diverse heterologous HIV trimers. B-C. BLI binding curves (B) and neutralization curves (C) of two representative mAbs, CH35-Apex1.08 and CH42-Apex2.01, isolated from protein-immunized rhesus macaques CH35 and CH42. These mAbs demonstrated binding and moderate neutralization breadth against the autologous Q23 panel, heterologous V2-apex-sensitive trimers, and the global HIV panel. Notably, both mAbs also neutralized chimpanzee Env-derived viruses, CAM13RRK and MT145K, which share the V2-apex bnAb site with HIV-1. D. Summary of neutralization IC50 values for V2-apex mAbs tested in panel B. Notably, CH35-Apex1.08 neutralized eight heterologous viruses, some with potent activity.

To compare the B cell responses of the protein and mRNA group animals, we used the same workflow to isolate V2-apex site-specific single B cells from week 14/16 LNs of six Q23-APEX-GT2 trimer mRNA-vaccinated group animals. Consistent with the serological data, the percentage of antigen- and epitope-specific B cells was lower in the mRNA group compared to the protein group (Figure S8B). Somatic hypermutation (SHM) analysis showed modest nucleotide divergence from the respective germline V genes in both the heavy (median = ∼4.5%) and light (median = 3.5%) chains for protein group and relatively lower for the mRNA group (Figure 4D, E), which is consistent with the early stages of affinity maturation of these B cell responses. Overall, protein immunized animals had superior antibody responses compared to the mRNA immunized group, which could have been due to the dose escalation prime regimen, the adjuvant and/or antigen display or suboptimal responses to the mRNA/LNPs when given subcutaneously rather than intramuscularly. We therefore focused on characterizing B cell and antibody responses in the protein immunized animals.

### Q23-APEX-GT2 induces long CDRH3 V2-apex bnAbs

We identified one to four long CDRH3 loop (22AA or longer) prototypic V2-apex bnAb-like lineages in week 14 or 16 LNs from each of the six Q23-APEX-GT2 trimer protein-vaccinated monkeys (Figure 4F). Since we only focused on week 14/16 LNs, it is conceivable that additional long CDRH3 lineages were present at the same or different time points. Each of the isolated lineages possessed D-gene (IGHD3-15, IGHD3-17 or IGHD3-18)-encoded anionic residues and predicted sulfated tyrosine motifs (e.g., EDDY, DDY, DDYDY, YDED, YEDD, DYY, YYD) in their CDRH3 paratopes (Figure 4F) known to enable interaction with the positively charged V2-apex core bnAb epitope^28,56,57,61^. These long CDRH3 B cells utilized a diverse set of heavy and light chain V and J genes, suggesting that the germline D-gene encoded motifs were likely the primary drivers of antigen-specific B cell selection.

Epitope mapping using BLI and virus neutralization showed that the long CDRH3 lineages bound to the Q23-APEX-GT2 trimer in an epitope-specific manner (Figure 4F). These mAbs also bound the WT Q23.17 trimer (Q23-APEX-GT1), demonstrating their ability to recognize native-like Env trimer configurations associated with the virus. Recognizing and maturing bnAb UCA precursors toward native-like trimer configurations is considered a key step in guiding bnAb precursors along the bnAb maturation pathway ^37,68^. The mAbs exhibited trimer-dependent binding (with no binding to monomeric Q23-APEX-GT2 gp120 (Figure S8C) and showed variable dependence on the N160 glycan (Figure S8D). While binding decreased for some mAbs with the N160-deficient Q23-APEX-GT2 trimer, others showed enhanced or unchanged binding (Figure S8D). Notably, many of these mAbs neutralized not only the autologous Q23-APEX-GT2-matched pseudovirus but also the WT Q23.17 virus in an epitope-dependent manner (Figure 4F).

We next investigated the BLI binding and neutralization of the long CDRH3 loop mAbs against multi-subtype tier-2 globally representative HIV-1 virus panels^43,69^. Notably, multiple mAbs from four monkeys exhibited cross-reactive binding while CH35-Apex1.08 and CH42-Apex2.01 also neutralized up to 8 heterologous tier-2 viruses including two chimpanzee SIV viruses that share V2-apex bnAb site with HIV-1 but are evolutionarily more distant than group M viruses (Figure 6A-D, S8E) ^40,70^. These findings mark the first demonstration of authentic V2-apex bnAb precursor induction and partial maturation into cross-neutralizing antibodies in an outbred monkey model using a germline-targeting immunogen. Thus, Q23-APEX-GT2 trimer immunization not only primed the authentic V2-apex bnAb precursors but also partially matured them along desired pathways for bnAb development.

**Figure 6.**
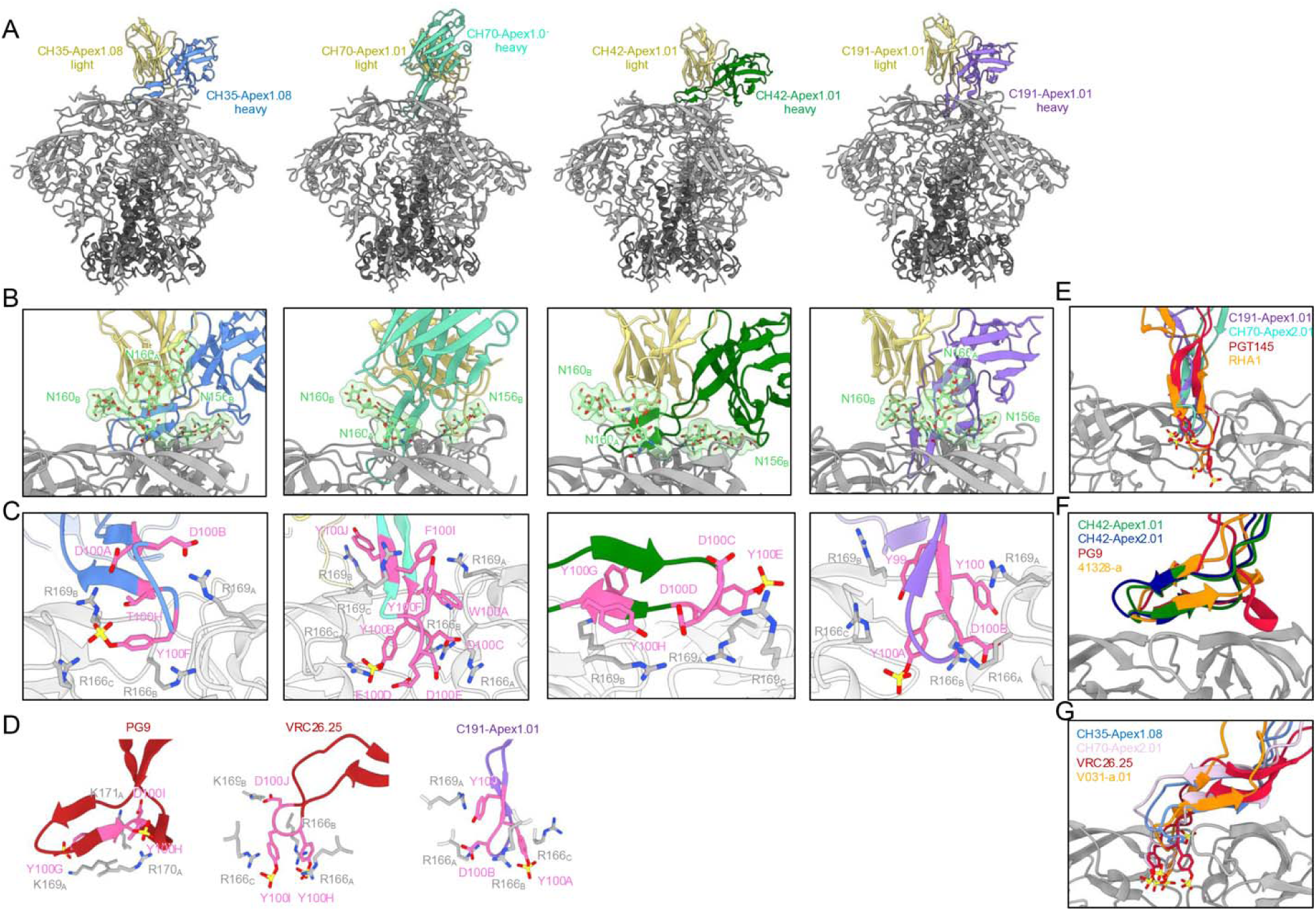
Cryo-EM structures of Q23-APEX-GT2-elicited antibodies reveal mimicry of canonical V2 apex bnAb recognition modes. See also Figures S9, S10, S11, and Dataset S1. A. Overall structure of Fab:envelope trimer complexes of V2-apex-targeted antibody lineages from four different Q23-APEX-GT2 immunized macaques determined by single-particle cryo-EM. All light chains are colored yellow while each heavy chain is colored by lineage. Envelope gp120 is shown in light gray and gp41 in dark gray. B. Expanded interface view of panel (A) to highlight Fab binding orientation and interactions with apical envelope glycans. Recognized glycans are shown in stick representation and colored green with transparent surfaces. Fab heavy and light chains are colored similarly to panel (A). C. Critical V2-apex gp120 contacts by HCDR3 germline-encoded D-gene residues. Fab heavy chains are colored by lineage similarly to panel (A), with paratope residues that are germline-encoded or conservatively mutated—somatic mutations that preserve the biochemical properties of the original germline residue—colored pink, shown in stick representation, and labeled by Kabat numbering. Sulfur atoms from sulfated tyrosine residues are colored yellow; nitrogen atoms are colored blue; and oxygen atoms are colored red. D. Structure and function of the D-gene germline-encoded three-residue YYD motif in human (PG9 and VRC26.25) and rhesus (CI91-Apex1.01) lineages. Fab HCDR3s from their respective complex structures are colored red (human) or purple (rhesus CI91), with the YYD motif colored pink, shown in stick representation, and labeled by Kabat numbering. Envelope V2 residues contacted by each respective YYD motif are shown with stick representation in isolation from the remainder of the gp120 structure. E-G. Structural superimpositions of Q23-APEX-GT2 vaccine-elicited V2-apex antibodies with mature human and rhesus V2-apex bnAbs using “combined-mode” VRC26-like (E), “needle” PGT145-like (F), and “axe” PG9-like (G) modes of recognition. Each structure is aligned by gp120 to compare V2 apex-bound Fab HCDR3 conformations and orientations. Mature human bnAb lineages are colored red and mature rhesus bnAb lineages are colored orange, with the vaccine-elicited antibodies colored similarly to panel (A).

An important question for targeted B cell approaches is what CDRH3 length and paratope features qualify as a bona fide V2-apex bnAb precursor. We observed that all B cells with CDRH3s of 22AA or longer bound as well as neutralized WT Q23 in a V2-apex bnAb epitope specific manner. However, while nearly all 23AA CDRH3 antibodies bound heterologous HIV trimers, the 22AA CDRH3-bearing V2-apex-targeted antibodies did not (Figure S8E). This pattern extended to neutralization, with only V2-apex-targeted antibodies with CDRH3s of 23AA or longer showing cross-neutralization of heterologous viruses. These findings suggest that a 23AA CDRH3 length may be the minimum requirement for a *bona-fide* V2-apex bnAb precursor ^57,61^. Whether 22AA or shorter CDRH3s can affinity mature into canonical V2-apex bnAbs remains to be investigated.

Based on these observations, we conclude that a *bona fide* V2-apex bnAb B cell precursor must meet the following criteria: i) a 23AA long CDRH3 loop with appropriate paratope features, ii) strand C epitope-dependent binding, iii) binding to both autologous WT and V2-apex-sensitive heterologous trimers, and iv) potential for autologous or heterologous virus neutralization. The last criterion (neutralization) may be a high bar, as these B cells are still early in their development with limited somatic mutations. Nevertheless, we believe that as *bona fide* bnAb precursors accumulate SHMs through vaccination, they should exhibit detectable neutralization of V2-apex-sensitive heterologous trimers, which would help differentiate them from strain-specific or non-bnAb B cell precursors. Based on these criteria, the Q23-APEX-GT2 trimer successfully induced *bona fide* V2-apex bnAb precursors in 4 out of 6 protein vaccinated monkeys in our study.

Overall, the B cell analysis was consistent with polyclonal plasma analysis, with both showing reproducible targeting of the V2-apex bnAb site. Thus, our findings provide proof-of-concept that authentic bnAb precursors can be primed using a germline-targeting vaccination approach and that some breadth can be gained without a complex prime-boost immunization protocol

### Structural analysis reveals shared features of elicited rhesus and human V2-apex bnAbs

Previous studies have shown three reproducible modes of HCDR3-dominated V2-apex epitope recognition: insertion of an extended “needle-like” HCDR3 directly into the trimer apex hole along the 3-fold axis ^41,55–58,61^; parallel or antiparallel main-chain hydrogen-bonding with the C-strand from a single protomer via an “axe-like” HCDR3 ^41,53,54,61^; and a “combined-mode” HCDR3 that simultaneously engages the C-strand through main-chain bonding and inserts the loop tip into the trimer hole ^56,61^. We have previously defined lineages utilizing these modes of V2 apex recognition as members of an “extended-class,” which are antibodies that utilize structurally similar paratopes derived from unique immunogenetic origins ^41^. Thus, we would propose that the one important criterion to define a bona fide V2-apex bnAb B cell precursor is: belonging to an extended-class characterized by one of three canonical modes of V2-apex bnAb recognition. To investigate whether the immunogenic and phenotypic properties of the Q23-APEX-GT2-elicited antibodies in fact recapitulated V2-apex bnAb recognition at the molecular level, we thus used single-particle cryo-EM to determine the structures of Fabs from six lineages (from four macaques: CH35, CH70, CH42 and CI91) in complex with Q23-APEX-GT2 trimer (Figure 6, S9, Dataset S1).

The cryo-EM structure of the CH35 macaque mAb CH35-Apex1.08 in complex with the Q23-APEX-GT2 trimer was resolved at 3.2 Å resolution. The structure revealed a 1:1 Fab:trimer stoichiometry with asymmetric binding along the 3-fold trimer axis (Figure 6A). CH35-Apex1.08 inserted between the N156 and N160 glycans on a single protomer while also engaging the N160 glycan from an adjacent protomer, leading to 52% of the total binding surface being mediated by apical glycans (Figure 6B, S10A). CH35-Apex1.08 utilized a combined-mode HCDR3 topology, similar to bnAbs VRC26.25 (human) and V031-a.01 (macaque) ^56,61^. It engaged the C-strand through three parallel main-chain hydrogen bonds while inserting an anionic loop tip—containing a sulfated tyrosine—into the trimer hole (Figure 6C, G, S10B) ^13,28,53–55,57^. This tyrosine sulfation, a common posttranslational modification in V2 apex bnAbs, facilitated interactions with core V2-apex residues R166, R168, and R169 from one or more Env protomers. Additionally, K171 on protomer B (K171_B_) was positioned between LCDR1 and LCDR3. Notably, IGHD3-15 D-gene germline-encoded residues played a key role in stabilizing interactions across all three Env protomers. D100A and D100B formed electrostatic interactions with residue R169 from promoter A (R169_A_) and residue R169 from promotor B (R169_B_), while sulfated Y100F engaged R169_A_ and residue R166 from promotor C (R166_C_) via electrostatic interactions and R166_B_ through cation-π interactions. The methyl group of T100H further stabilized binding by engaging the aliphatic chain of R169_B_. Unlike human bnAb VRC26.25, which inserts a second sulfated tyrosine into the trimer hole, rhesus bnAbs often substitute it with an anionic residue ^61^. In line with this, CH35-Apex1.08 acquired a D100G somatic mutation (Tyr to Asp), allowing electrostatic contacts with R166_A_ and R169_C_, effectively completing recognition of the cationic lining of the trimer hole (Figure S10C).

The cryo-EM structures of the CH70 macaque mAbs CH70-Apex2.01 and CH70-Apex1.01, resolved at 3.8 Å and 3.0 Å respectively, revealed distinct modes of V2-apex recognition (Figure 6A, S9). CH70-Apex2.01 bound near the trimer 3-fold axis with a 1:1 Fab:trimer stoichiometry, engaging three apical glycans, though these interactions accounted for only 18% of the total binding surface (Figure 6B, S10A). The antibody primarily engaged Env through an extended, needle-like HCDR3 that inserted directly into the trimer hole, resembling the recognition mode of bnAbs PGT145 (human) and RHA1 (macaque) (Figure 6C, E) ^55,57^. Extensive contacts with core V2-apex residues were mediated by paratope residues that were either IGHD3-15 germline-encoded or conservatively mutated—somatic mutations that preserve the biochemical properties of the original germline residue (e.g., Asp to Glu maintaining negative charge or Tyr to Trp retaining aromaticity) (Figure 6C). Aromatic residues Y100F and F100I recognized R169_C_ via hydrogen bonding and cation-π interactions, while W100A and Y100J engaged R169_A_ and R169_B_ from other protomers. Further down into the trimer hole, the anionic HCDR3 tip— comprising sulfated Y100B, D100C, E100D, and D100E—formed salt bridges with R166 from all three protomers. Antibody CH70-Apex1.01 also bound Env with a 1:1 Fab:trimer stoichiometry but approached the three-fold axis at a more asymmetric angle (Figure S9). Similar to CH70-Apex2.01, it engaged three apical glycans but contributed a greater fraction (36%) of the total binding surface (Figure S10A). CH70-Apex1.01 utilized a combined-mode HCDR3 topology, inserting two germline-encoded sulfated tyrosines into the trimer hole while simultaneously forming three main-chain hydrogen bonds with the C-strand (Figure 6G, S10B). Similar to human bnAb VRC26.25, these sulfated tyrosines formed electrostatic contacts with R166 from all three Env protomers. Additional interactions included D100C forming a salt bridge with R169_C_ and Y100D stabilizing the extended aliphatic chain of R169_A_ (Figure S9C, left).

The cryo-EM structures of CH42 macaque mAbs CH42-Apex1.01 and CH42-Apex2.01, resolved at 2.9 Å and 3.1 Å respectively, showed both antibodies binding with 1:1 Fab:trimer stoichiometry through an axe-like HCDR3, which formed parallel strand bonding with the C-strand (Figure 6A, F, S9, S10A). This recognition mode mirrors that of bnAbs PG9 (human) and 41328-a.01 (macaque) ^53,61^. CH42-Apex1.01 inserted between the N156 and N160 glycans on a single protomer while also engaging the N160 glycan from an adjacent protomer, resulting in 54% of the Fab binding surface being mediated by apical glycans (Figure 6B, S10A). While three parallel main-chain hydrogen bonds were formed with the C-strand of protomer B, the HCDR3 of CH42-Apex1.01 extended across the trimer hole, allowing IGHD3-15 germline-encoded residues to interact with core V2-apex residues from all three protomers (Figure 6C). Specifically, D100C and D100D were positioned near R169_C_ and R169_B_, respectively, suggesting potential electrostatic interactions. The sulfated Y100E formed a salt bridge with R169_C_ while simultaneously stabilizing the extended aliphatic chain of R169_A_. Additionally, Y100G engaged K168_B_ via cation-π interactions, and Y100H interacted with R169_B_ through both aliphatic chain stabilization and hydrogen bonding with the N-terminal amine. Antibody CH42-Apex2.01 similarly bound asymmetrically to the trimer 3-fold axis, engaging three apical glycans that accounted for 46% of the Fab interactive surface (Figure S9, S10A). Its interactions included four main-chain hydrogen bonds with the C-strand, three through parallel strand bonding and one via N100I carboxyamide sidechain with K171 backbone amide. Notably, cryo-EM density revealed three posttranslational sulfation modifications on IGHD3-15 germline-encoded residues Y100D, Y100F, and Y100H. These modifications collectively facilitated electrostatic interactions with R169 from all three protomers and R166 from protomers A and C (Figure S9C, right). Additional contacts included salt bridges with R169_A_ (via D100B) and R166_A_ (via D100E), plus multiple light chain interactions with K171_B_ mediated by LCDR1 and LCDR3 (Figure S10D).

The cryo-EM structure of CI91 macaque mAb CI91-Apex1.01 complex was resolved at 3.0 Å, showing a 1:1 Fab:trimer stoichiometry with binding near the three-fold trimer axis (Figure 6A). Similar to other Q23-APEX-GT2-elicited antibodies, CI91-Apex1.01 recognized three apical glycans—N160 and N156 from one protomer and N160 from an adjacent protomer—accounting for 30% of the total interactive surface (Figure 6B, S10A). CI91-Apex1.01 primarily engaged Env through its HCDR3, which adopted an extended needle-like conformation with two sulfated tyrosines flanking the loop tip that inserted directly into the trimer apex hole. This binding mode closely resembled that of mature human and rhesus bnAbs in the PGT145-extended class (Figure 6E). CI91-Apex1.01 recognized R166 and R169 from all three protomers, with interactions largely mediated by IGHD3-18 D-gene germline-encoded residues forming the descending HCDR3 β-strand and part of the loop tip. Specifically, Y100 was positioned between R166_B_ and R169_A_, stabilizing their extended aliphatic chains. Sulfated Y100A engaged R166_B_ and R166_C_ through electrostatic interactions and also formed a hydrogen bond with the backbone amide of R166_B_. Additionally, D100B was inserted between R166_A_ and R166_B_ (Figure 6E). The second sulfated tyrosine, Y100F, played a critical role in the paratope by recognizing both R166_C_ and R169_B_ (Figure S10E). Structural alignment of CI91-Apex1.01 with CH70-Apex2.01, another needle-like V2 apex bnAb precursor, showed that both Q23-APEX-GT2-induced antibodies aligned their HCDR3s with those of mature bnAbs VRC26.25 and RHA1— the first human and rhesus bnAbs described in this extended class. However, unlike the mature bnAbs, CI91-Apex1.01 and CH70-Apex2.01 did not penetrate as deeply into the trimer hole (Figure 6E). Instead, the sulfated HCDR3 loop tips overlapped with the mature rhesus V2 apex bnAb 44715-a (Figure S10F) ^61^.

CI91-Apex1.01 was notable for its utilization of IGHD3-18 germline D-gene, in contrast to IGHD3-15, which has been consistently used by all other structurally characterized rhesus V2-apex bnAbs and their precursors, both in this study and previous reports ^57,61,71^. While IGHD3-15 encodes a rhesus-specific EDDY motif, IGHD3-18 encodes a three-residue YYD motif, which is also found in human IGHD3-3. This motif is shared by three human V2-apex bnAb lineages— PG9, PCT64, and VRC26.25—all of which originate from IGHD3-3 ^8,28,46,50^. The mature bnAbs PG9 and VRC26.25 retain the YYD motif to mediate electrostatic interactions with core V2-apex epitope residues, although through unique structural mechanisms (Figure 6D). CI91-Apex1.01 represents a third, structurally divergent conformation of this conserved motif while maintaining its functional role in interacting with cationic V2-apex residues. Additionally, it retains the characteristic posttranslational sulfation modification on the IGHD3-3/IGHD3-18 germline-encoded tyrosines.

Overall, this structural analysis provides explicit molecular evidence for the ability of Q23-APEX-GT2 to induce bona fide V2 apex bnAb precursors in multiple outbred macaques and reveals how germline-encoded D-gene residues compose the paratope recognizing critical V2 apex contact residues, most commonly at positions 166 and 169. These precursors fell into previously defined bnAb extended-classes which recapitulated all three modes of mature V2 apex bnAb recognition, with two modes of recognition being observed even within a single macaque, demonstrating that Q23-APEX-GT2 can prime a diverse pool of precursors with distinct structural features. Notably, all vaccine-elicited rhesus antibodies recognized the engineered germline-binding-enhancing T132R modification via salt bridges mediated by one or two anionic residues, most commonly (5/6 lineages) through the light chain (Figure S11). The result suggests that the T132R modification likely improved the vaccine priming efficacy of Q23-APEX-GT2 by selectively engaging V2-apex precursors through direct epitope-paratope interactions.

### Lineage tracing of long CDRH3 V2-apex bnAb B cell responses by bulk NGS

To track the population-level dynamics of long CDRH3 (23 amino acids or longer) B cell responses following Q23-APEX-GT2 immunizations, as well as the expansion and longitudinal development of isolated V2-apex epitope-specific B cells, we performed next-generation sequencing (NGS) of lymph node B cells at weeks 0, 4, 8, and 12 using a bulk IgG and IgM immunoglobulin amplification approach. Prior to immunization, animals in both the protein and mRNA vaccine groups exhibited a typical Gaussian distribution of B cell CDRH3 loop lengths with a peak at 14AA (Figure S12). Consistent with the rhesus baseline repertoire, a small fraction of long CDRH3 B cells (range: 0.15 −0.55 %) was observed across all animals, predominantly of the IgM isotype (Figure 7A, S12). Post-immunization, B cell repertoires were highly enriched for longer CDRH3s, with median lengths increasing from 14 amino acids (baseline in monkeys) to 16–18 amino acids. This shift was largely driven by B cells with CDRH3s of 23 amino acids or longer (Figure S12). At week 4, a more rapid enrichment of long CDRH3 B cells was observed in the protein-vaccinated group compared to the mRNA group (Figure S12). However, by week 12 (two weeks post-second immunization), the two groups were comparable. The shift toward longer CDRH3s was primarily contributed by class-switched IgG B cells, a trend that was consistent across both vaccine groups (Figure S12).

**Figure 7.**
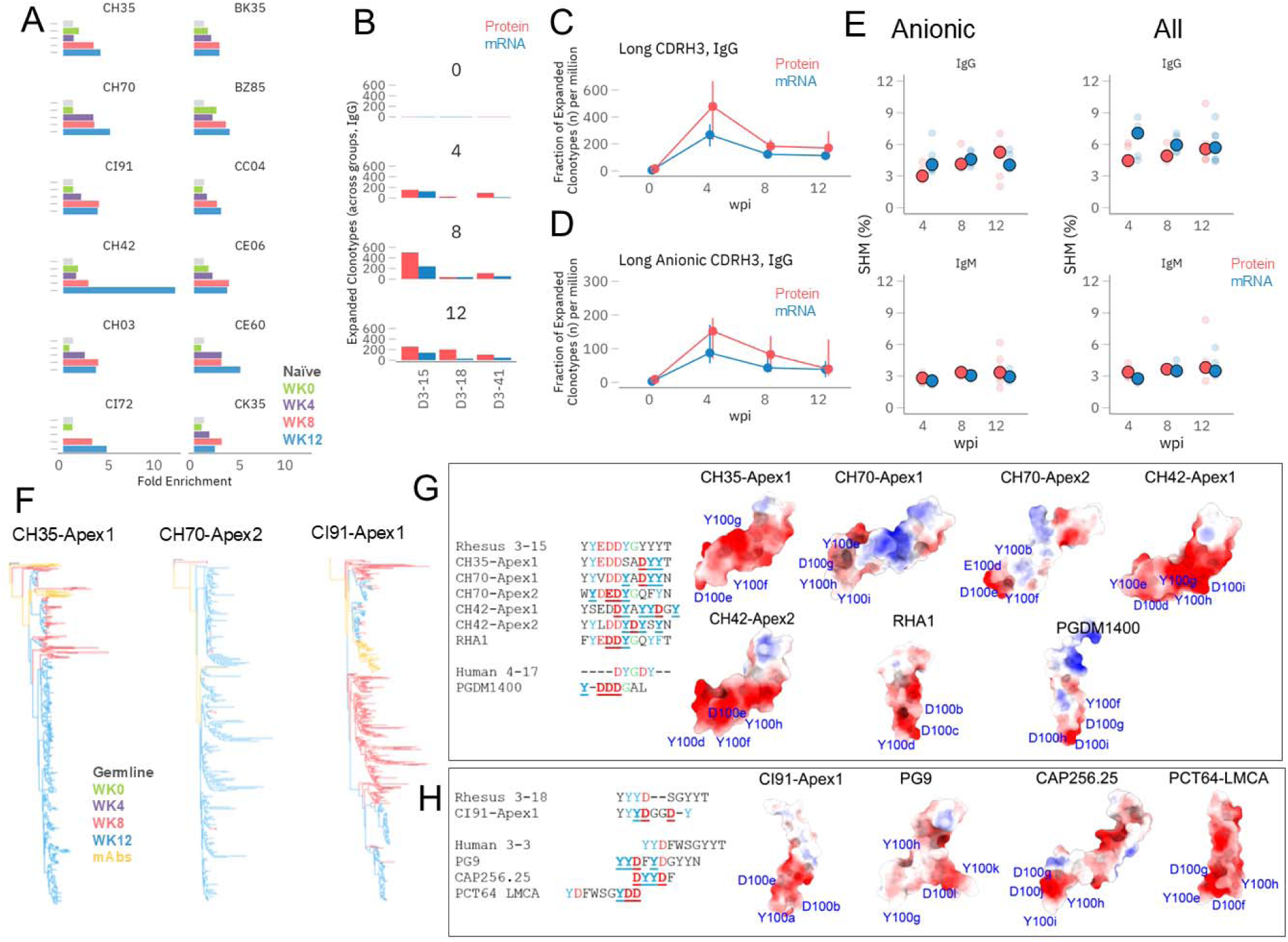
B cell lineage tracing of long CDRH3 V2-apex mAbs in Q23-APEX-GT2 trimer protein and mRNA immunized macaques by bulk Next-Generation Sequencing (NGS). See also Figures S12, S13 and S14. A. Longitudinal enrichment of long CDRH3 B-cells (≥23 AA) for individual RMs in Q23-APEX-GT2 protein and mRNA vaccinated animals. LN samples from time points, weeks 0, 4, 8 and 12 were analysed a and compared with baseline naïve macaque repertoires. B. Longitudinal enrichment of anionic D genes within the expanded clonotype IgG compartment. Protein is colored red, mRNA is colored blue. Certain D-genes show enrichment as the immunization progresses indicating antigen driven selection. C. Longitudinal enrichment of all long CDRH3s within expanded clonotypes in Q23-APEX-GT2 immunized animals. D. Longitudinal enrichment of anionic long CDRH3s within expanded clonotypes in Q23-APEX-GT2 immunized animals. E. Longitudinal increment in SHM levels within the IgG compartment of Q23-APEX-GT2 trimer protein-immunized animals (top half) within B cells bearing anionic CDRH3s. F. Phylogenetic trees show longitudinal development V2-apex mAb lineage in animals CH35, CH70, and CI91. The week 14 isolated mAbs are clustered with B cell sequences from weeks 0, 4, 8, and 12 derived from bulk NGS. G-H. Two rhesus germline D-gene solutions (IGHD3-15 and IGHD3-18) share commonalities with human germline D-gene solutions (IGHD4-17 and IGHD3-3). Both incorporate anionic CDRH3 residues and sulfated tyrosine motifs, which are critical for V2-apex bnAb site recognition. While the rhesus IGHD3-15-encoded EDDY motif resembles the human IGHD4-17-encoded DY motif (G), the rhesus IGHD3-18-encoded YYD motif is identical to the human IGHD3-3-encoded YYD motif (H). Q23-APEX-GT2 successfully induced V2-apex bnAb precursors in rhesus macaques carrying either of these motifs. Alignments of CDRH3 amino acid sequences from Q23-APEX-GT2 vaccine-elicited V2-apex mAbs are compared with corresponding rhesus and human germline D-gene sequences, highlighting the anionic motifs involved in V2-apex epitope recognition. Structural analysis of the CDRH3 loops of these mAbs reveals that the apical positioning of this germline-encoded motif is essential for recognizing the V2-apex bnAb site.

One limitation of bulk NGS is that each B cell can contribute multiple copies of the same Ig transcript, potentially skewing the analysis. This issue is particularly pronounced for plasmablasts or plasma B cells, which express very high levels of Ig mRNA transcripts and could misrepresent the magnitude of B cell responses. To address this bias, we analyzed long CDRH3 loop IgG B cells focusing only on expanded clones (>3 clonal members). This approach revealed a similar trend of long CDRH3 B cell expansions, with both vaccination groups appearing comparable (Figure 7A). However, compared to the mRNA group, protein-vaccinated animals exhibited a substantially higher number of long CDRH3 B cell clonotypes with appropriate anionic paratope motifs characteristic of V2-apex bnAbs (Figure 7B-D), consistent with earlier findings indicating superior responses in the protein group. Long CDRH3s were highly enriched in IGHD3-15, but certain other rhesus germline D genes, such as IGHD3-18 and IGHD3-41, also contributed to the expanded long CDRH3 lineages (Figure 7B, S13A, C).

We investigated the levels of somatic hypermutation (SHM) in the VH region of IgM and IgG sequences at weeks 4, 8, and 12 post-immunizations (Figure 7E), which ranged from ∼3-7% (Figure 7E). While SHM levels in IgM B cells remained largely unchanged over time across both immunization groups, SHMs in class-switched IgG B cells, particularly in long CDRH3 B cells, increased over time in the protein-vaccinated group. We also observed the elevated prevalence of long CDRH3 B cells with sulfated tyrosine CDRH3-motifs (Figure S13). These findings suggest that long CDRH3 B cell lineages were under strong antigen-driven selection, consistent with efficient priming of long CDRH3 bnAb B cell lineages.

To track the lineage development of the prototype V2-apex site-specific mAbs isolated at week 14 (shown in Figure 4F), we surveyed their corresponding lineages in the longitudinal NGS data. Most of these site-specific lineages were expanded, with a few exceptions (Figure 7F, S13). NGS lineage tracing revealed that most prototype long CDRH3-expanded lineages were likely activated by the immunization prime. These lineages became detectable by week 8 but were mostly not observed at the week 4 time point. This observation may have been influenced by B cell expansion size and trafficking, as the analysis was based on LN biopsies, with different LNs sampled at various time points. Overall, the data demonstrate that the Q23-APEX-GT2 trimer efficiently primed rare prototype V2-apex long CDRH3 bnAb B cells after the first vaccination and subsequent homologous boost, in most cases, facilitated robust expansion and further evolution of these bnAb lineages.

NGS analysis revealed that, compared to the mRNA group, the protein group animal exhibited a greater number of prototype V2-apex-like expanded long CDRH3 lineages. The magnitude of these expansions significantly increased following the boost immunization, and we identified more than 10 long CDRH3 B cell lineages exhibiting V2-apex bnAb-like features in some animals. Overall, NGS identified a relatively larger number of expanded prototype V2-apex bnAb-like long CDRH3 B cell precursor clones compared to those found through antigen epitope-specific mAb isolation. To determine whether pre-existing IgM or IgG V2-apex-like long CDRH3 B cell precursor frequencies influenced the elicitation and expansion of these lineages, we analyzed pre-bleed lymph node LN NGS data. The analysis revealed comparable levels of IgM- and IgG-encoded long CDRH3 V2-apex bnAb-like B cell precursors across all animals (Figure S13D), suggesting that pre-existing precursors had minimal influence on the observed differences across animals.

### Rhesus V2-apex long CDRH3 B cell repertoires: lessons for human vaccination

To gain deeper molecular insight into the Q23-APEX-GT2-elicited rhesus V2-apex bnAbs and their relevance to targeted human vaccine strategies, we closely examined the germline D-gene–encoded paratope motifs. Our analysis revealed that, while Q23-APEX-GT2 efficiently engages rhesus IGHD3-15–encoded bnAb lineages bearing the CDRH3 EDDY motif, it also successfully engages a IGHD3-18–encoded bnAb in one rhesus macaque (CI91), which incorporates the YYD motif (Figure 7G, H). This finding is unique to our study, as all previously reported rhesus V2-apex bnAbs are encoded exclusively by the IGHD3-15 gene ^57,61^. This underscores the potential of Q23-APEX-GT2 to engage diverse but convergent epitope-recognizing paratope solutions. Interestingly, the rhesus IGHD3-15–encoded EDDY paratope closely mirrors the human IGHD4-17–encoded DY motif found in one of the most potent and broadly neutralizing human bnAb prototypes, PGDM1400 ^72^. In contrast, the rhesus IGHD3-18–encoded YYD paratope resembles the human IGHD3-3-encoded YYD motif observed in three human V2-apex bnAb prototypes: PG9, CAP256, and PCT64 ^8,28,46,50^. While a variety of D-gene–encoded CDRH3 anionic motifs are present in both infection- and vaccine-elicited human and rhesus V2-apex bnAb prototypes, the apical positioning of the CDRH3 motif—allowing interaction with the buried, positively charged strand C - is critical for binding, as supported by structural studies (Figure 7H).

To assess the relevance of V2-apex antibody germline features observed in rhesus macaques for human vaccine strategies, we analyzed the frequency of key bnAb-encoding CDRH3 motifs in the human naïve B cell repertoire ^60^. The median occurrences (per million B cells) of these motifs at apical positions in long CDRH3s (≥23 amino acids) were: YYD (895, range 640–2161), DY (759, range 460–1225), DDY (19, range 11–35), (E|D)(E|D)Y (42, range 22–65), and (E|D)DDY (9, range 4–17) (Figure S14, Supplementary Table). These data reveal a clear trend: as anionic motif complexity increases, its frequency in the human repertoire declines. Simple motifs like DY and YYD are common, whereas DDY is an order of magnitude rarer, and (E|D)DDY is the least frequent. This suggests that while the immunogenetic features engaged by Q23-APEX-GT2 immunization in macaques exist in humans, they are significantly less prevalent. Their rarity implies that effective precursor priming strategies may be needed to expand and engage these bnAb B cell precursors. Overall, our study demonstrates that rhesus macaques are a useful model for inducing V2-apex bnAbs with features in common with humans, although there are potentially important differences.

## DISCUSSION

Molecular vaccine design offers significant promise in overcoming the challenges of inducing protective bnAbs against HIV ^18,21,24^. Our approach leverages molecular insights of HIV bnAbs to guide the development of immunogens that target specific bnAb precursors and direct their maturation. In this study, we applied antibody-guided structure-based design to develop a germline-targeted (GT) trimer immunogen targeting the V2-apex bnAb site, a key broadly neutralizing epitope on HIV Env. Immunization of outbred macaques with our engineered Q23-APEX-GT2 immunogen successfully expanded and partially matured HIV V2-apex bnAb precursors, demonstrating for the first time that cross-neutralizing antibodies can be induced by a germline-targeting immunogen in an outbred macaque model. These findings highlight the V2-apex as a particularly promising target for vaccination and suggest that bnAb induction at this site may be achievable including in humans through a relatively straightforward pathway.

Priming rare HIV bnAb precursors is a critical step in bnAb induction ^73–75^. The V2-apex bnAb site is a highly promising vaccine target recognized by some of the most potent and broad mAbs; however, priming its precursors remains challenging due to the rarity of long CDRH3 precursors with necessary structural and biochemical features ^13,28,59^. To address this, we developed a priming immunogen for the V2-apex bnAb site based on the Q23.17 HIV Env, which has consistently induced V2-apex bnAbs in rhesus macaques in a SHIV infection model ^61^. The Q23 GT-immunogen was engineered to engage diverse V2-apex bnAb precursors, to increase the likelihood of priming rare bnAb B cells *in vivo*. Analyses of Q23 GT-trimer elicited polyclonal immune sera demonstrated strong immunofocusing to the V2-apex site, with responses dependent on the N160 glycan and strand C region residues, key components of the V2-apex bnAb site. This strong antibody immunofocusing is noteworthy, given the complex antigenic landscape of the HIV glycoprotein surface, making the Q23-APEX-GT2 trimer an excellent candidate for priming bnAb precursors.

Antigen-specific B cell analysis revealed the induction of V2-apex-specific long CDRH3 antibodies that bound a genetically and antigenically diverse panel of HIV Envs. Despite using only a single homologous prime and boost immunization, some of these antibodies affinity matured to exhibit heterologous tier-2 HIV neutralization. The vaccine-induced neutralizing antibodies displayed structural and genetic features resembling human V2-apex bnAbs, encompassing all three known CDRH3 configurations ^61,76^. This study is the first to show that GT-trimer immunization alone can successfully prime authentic V2-apex bnAb B cell precursors and drive their partial maturation into cross-neutralizing antibodies in outbred macaques. In contrast, germline-targeting efforts for other HIV Env sites have never achieved neutralization or even cross-reactive binding to native-like HIV trimers in outbred monkeys ^31,32^. One potential reason is that Q23-APEX-GT2 in our study was minimally modified at the V2-apex bnAb site, preserving a near-native trimer configuration while enhancing affinity for diverse prototype rhesus and human bnAb UCAs. This may enable affinity maturation to be driven by a native-like trimer, selectively shaping primed BCR configurations for bnAb development. This may define V2-apex bnAb germline-targeting strategies, where preserving the native-like trimer architecture at the bnAb epitope—despite engineering modifications—is essential for engaging long bnAb B cell precursors that rely on trimer-dependent recognition. Overall, the findings show that V2-apex bnAbs, which heavily rely on long CDRH3-encoded residues for epitope recognition, can mature into early bnAbs with minimal diversification in the boost regimen, a result that aligns with observations of vaccine-induced ultra-long CDRH3 bovine HIV bnAbs ^77,78^.

An important consideration in translating rhesus macaque-based V2-apex bnAb vaccine strategies to humans is the potential difference in germline D-gene features that affect the initial engagement of germline-targeting immunogens with bnAb B cell precursors. While rhesus and human V2-apex bnAbs share immunogenetic and structural similarities, a key distinction lies in their D-gene-encoded anionic motifs, which influence bnAb precursor recruitment and maturation. In rhesus, the IGHD3-15 germline D-gene encodes a highly anionic EDDY motif, facilitating electrostatic interactions with the positively charged V2-apex region of the HIV Env trimer. This feature is critical for early B cell engagement and subsequent affinity maturation toward bnAbs. The most common human V2-apex bnAb germline D-gene motif, found in prototypic bnAbs PG9, CAP256, and PCT64, is the IGHD3-3-encoded YYD motif, which can also originate from human germline D-genes, IGHD3-9, IGHD3-16, and IGHD3-22 ^28,60,79^. Additionally, human V2-apex bnAb precursors can encode rhesus-equivalent DY (encoded by IGHD4-17) or DDY (likely generated through secondary diversification) motifs. Compared to rhesus, all three human CDRH3-paratope solutions contain fewer anionic residues in their native configuration, which may limit their recruitment and evolution. However, the rhesus IGHD3-18 germline D-gene incorporates a YYD motif, and Q23-APEX-GT2 successfully engaged a bnAb precursor bearing this motif in our monkey trial. This observation suggests that immunogens designed to engage YYD-bearing rhesus bnAb precursors could expand the pool of potential bnAb-precursor lineages in humans, improving the translation of vaccine strategies to a broader class of human V2-apex bnAbs. Irrespective of the differences, the rhesus macaque offers the opportunity to test a full sequential immunization strategy – from precursor priming to bnAb maturation and polishing – and establish proof-of-principle for the strategy in an outbred animal model.

While our GT-trimer effectively activated V2-apex bnAb precursors, variability in priming efficiency across animals underscores the need for further optimization to consistently and robustly engage diverse precursors. We observed that GT-priming was sufficient to induce early cross-neutralizing antibodies with some neutralization breadth, necessitating boost strategies to enhance the breadth and potency of these antibodies. Q23 Env lineage-based boosts, with minimally engineered V2 C-strand, may effectively broaden V2-apex bnAb responses, as strand C diversity alone has been shown to be sufficient for V2-apex bnAb maturation in macaques and humans ^8,46,52,57,61,62^. This Env lineage-based approach is likely to reduce *de novo* B cell responses seen with heterologous boosts at secondary immunization ^80,81^, though it may face challenges from circulating antibody feedback effects ^82–84^. In our priming experiments, we did not observe extremely high antibody titers that may interfere with the boosting immunogens. Furthermore, it is important to note that heterologous boosts will reduce effects of antibody feedback ^82^, as the affinity of the existing antibodies will be reduced as a result of the epitope structural changes introduced in the immunogen. The optimal affinity distance between successive immunogens is also crucial for efficient bnAb memory B cell (MBC) recall and maturation ^26,34,52,85,86^. A key challenge for MBC recall is limited MBC participation in secondary GCs, often due to differentiation into plasma cells or recruitment of *de novo* primed naïve B cell responses during boosts ^87–89^. Therefore, in multi-stage HIV vaccination strategies, preserving the bnAb MBC pool at each immunization step is critical for their maturation and eventually differentiation into plasma cells at the final boost to induce durable antibody titers.

The overall goal of HIV vaccination is to elicit broad serum neutralization targeting multiple bnAb specificities. Our study demonstrates that bnAb induction through vaccination is feasible and provides a framework for targeting CDRH3 germline-encoded features—an approach that could be extended to other Env bnAb sites on the surface glycoproteins of highly evasive viruses. The V2-apex bnAb site is particularly suited for HIV vaccine targeting, offering a potentially simpler immunization solution compared to other Env bnAb sites. By overcoming challenges like V2-apex rare precursor activation and showing the potential of GT vaccines to coax the antibody response a considerable distance along a desirable maturation pathway enhances optimism on the delivery of an effective HIV vaccine.

### Limitations of the study

While our GT-immunogen successfully primed authentic V2-apex bnAb precursor B cells and elicited cross-neutralizing antibodies, the breadth and potency of these monoclonal antibodies require enhancement through boost immunization strategies. This remains a key limitation and an area for further study, with the ultimate goal of inducing broad, potent, and durable serum antibody responses that provide protection against diverse HIV isolates. Another limitation is the potential differences in the B cell repertoires between humans and rhesus macaques, which could influence B cell and antibody responses. These differences may impact the translation of vaccine efficacy observed in macaques to human studies, making this an important area of ongoing investigation.

## ACKNOWLEDGEMENTS

We thank the staff at Bioqual for exceptional care and assistance with nonhuman primates. This work was supported by National Institutes of Health grants R61 AI 161818 (R.A., G.M.S.), R01 AI 167716 (R.A.), P01 AI 177683 (R.A., B.B.), R01 AI 160607 and AI165080 (G.M.S.), R37 AI150590 (B.H.H.), the University of Pennsylvania Center for AIDS Research (P30 AI 045008), the National Institute of Allergy and Infectious Diseases (NIAID) Consortium for HIV/AIDS Vaccine Development (CHAVD; UM1 AI144371, UM1AI144462) (A.B.W., M.C., B.B., and D.R.B.), the Bill and Melinda Gates Foundation through the Collaboration for AIDS Vaccine Discovery grants INV041767 and INV064777 (G.M.S., and R.A.) and INV-070116 (M.C., J.D.A.). We thank Carlos Diazgranados and Pervin Anklesaria from Gates Foundation for their support and helpful discussions.

## AUTHOR CONTRIBUTIONS

N.M., B.L., R.S.R., A.G., S.C., D.R.B, F.D.B, P.D.K., G.M.S and R.A. conceived and designed experiments. N.M., B.L., S.C., X.L. A.L.V., G.A., R.R.C., A.S., S.P., C.M., K.D., T.C., R.N. prepared immunogens, isolated mAbs, performed neutralization and binding assays. N.M., B.L., X.L., Y.Z., R.H., F.B.R., S.P., A.N.S., A.S. performed B cell repertoire sequencing and analysis. N.M., B.L., S.C., X.L., A.L.V, G.G., Y.Z., M.K., performed single B cell sorting and B cell sequence analysis. N.M., B.L., S.C., X.L., A.L.V., G.A., G.G., P.O., K.A., Y.Z., Y.Z., M.K., L.T., S.A. cloned, expressed, purified and tested monoclonal antibodies and trimers and probes for B cell sorting. A.G., R.A.R., F.D.B., conducted the mouse studies. M.M.L. F.B.R., B.H.H., G.M.S. led NHP studies. E.B.A., K.K.M., and D.J.I., synthesized and provided SMNP adjuvant. J.D.A., N.E.J., M.C. performed glycan analysis. X.L., W.L. G.O., A.B.W., K.N.R., L.v.d.M. generated and analyzed EMPEM data. S.F., A.C., S.H. provided mRNA. R.S.R., L.S., P.D.K. performed cryo-EM high-resolution structures of antibodies in complex with trimers and structural analysis. N.M., B.L., F.B.R., Y.Z., J.H., B.B. analyzed repertoire and bulk cell NGS data. N.M., B.L., R.S.R., A.G., F.D.B., P.D.K., D.R.B., G.M.S. and R.A. wrote the manuscript with input from all listed authors.

## DECLERATION OF INTERST

R.A., G.M.S., R.N., X.L., B.L., R.R.C., Y.Z., K.A., N.M., S.C., G.A., D.R.B., and W.H. are listed as inventors on pending patent applications jointly filed by the University of Pennsylvania and Scripps Research, related to the HIV-1 immunogens described in this study. All other authors declare no competing interests.

## STAR METHODS

## CELL LINES AND EXPERIMENTAL MODELS

### Cell Lines

Expi293F cells (Gibco, Cat# A14527) were cultured in Expi293 Expression Medium (Gibco, Cat# A1435101) at 37°C with 8% CO₂ in a 125 rpm shaker. HEK293F cells (Gibco, Cat# A14527) were cultured in Freestyle medium (Gibco, Cat# 12338-018) at 37°C with 8% CO₂ in a 125 rpm shaker. HEK293T cells (ATCC, Cat# CRL-3216) were maintained in Dulbecco’s Modified Eagle Medium (DMEM) (Corning, Cat# 10-017-CV) supplemented with 10% heat-inactivated fetal bovine serum (FBS) (Thermo Fisher, Cat# MT35016CV), 4 mM L-glutamine (Corning, Cat# 25-005-CI), and 1% penicillin-streptomycin (P/S) (Corning, Cat# 30-002-CI) at 37°C in a 5% CO₂ incubator. TZM-bl 931 cells (NIH AIDS Reagents Program) were used for the pseudovirus neutralization assay as previously described.

### Animal models

#### Mice

Previously described PCT64^LMCA^ knock-in (KI) mouse^42^ derived B cell was used for immunization experiments to investigate the B cell priming efficiency of Q23-APEX-GT2 trimer as a protein and mRNA lipid nanoparticle. 8-12 weeks old B6.SJL-Ptprca Pepcb/BoyJ (CD45.1 mice) mice were used for B cell adoptive transfer/immunization experiments.

### Indian rhesus macaques

All 12 Indian Rhesus macaques used in this study were housed at Bioqual, Inc. (Rockville, MD) in compliance with the guidelines set by the Association for Assessment and Accreditation of Laboratory Animal Care (AAALAC). All experimental procedures were approved by the Institutional Animal Care and Use Committees (IACUC) of the University of Pennsylvania (protocol 807492) and Bioqual (protocol 24-072). Macaques were sedated for blood collection and received care in accordance with AAALAC guidelines and best practice standards.

### Immunogen Design

To evaluate the interaction of V2-apex broadly neutralizing antibody (bnAb) precursors with key V1V2 sites, a series of site-directed mutagenesis (SDM) experiments were performed on the base germline-targeting trimer construct (Q23-APEX-GT1). Mutagenesis targeted residues within the V1V2 region (positions 130, 132, 158, 167–174) that were identified from structures of rhesus and human V2-apex bnAbs in complex with HIV trimers as potential hotspots for enhanced bnAb precursor engagement. Mutagenesis was carried out using the NEB Q5 site-directed mutagenesis kit (New England Biolabs, cat #M0494S) following the manufacturer’s protocol, and successful incorporation of mutations was verified through sequencing analysis (Eton Bioscience, San Diego, CA).

### Stabilized Env Expression and Purification

Plasmids encoding the Env trimers were transfected into HEK293F cells using PEI-MAX 40K transfection reagent (Kyfora, cat# 24765-1). The cells were incubated, and four days post-transfection, the supernatants containing the expressed trimers were collected. Purification was carried out using affinity chromatography with either agarose-bound Galanthus nivalis lectin (GNL) (Vector Labs, cat #AL-1243-5) or TOYOPEARL AF-Tresyl-650M beads (TOSOH, cat #0014472) conjugated to the PGT145 broadly neutralizing antibody (bnAb). To ensure further purification, the eluates were subjected to size-exclusion chromatography (SEC) using a Superdex 200 increase 10/300 GL column (GE Healthcare, cat #GE28-9909-44) in PBS. SEC fractions corresponding to the trimer peak were pooled together and utilized for ELISA and BLI binding studies, immunizations and as baits for antigen or epitope specific single B cell flow cytometry sorting experiments. Monomeric Q23 gp120 was purified by GNL affinity followed by SEC segregation and selection of the protein fractions corresponding to the gp120 monomer peak.

### Site-specific glycan analysis

To determine the glycosylation of the Q23_SCT27_GT2.V1 trimer, 100µg of protein was denatured for 1-hour in 50mM Tris/HCL, pH 8.0, containing 6M of urea and 5mM dithiothreitol (DTT). The sample was then incubated in the dark for 1-hour with 20mM iodoacetamide (IAA) to alkylate the protein. To remove the residual IAA, 20mM DTT was added, and the sample was incubated for an additional 1-hour period. Following buffer-exchange into 50mM Tris/HCL, pH 8.0 using Vivaspin columns (10kDa), the proteins were split into three aliquots, each containing approximately 33µg of protein to allow for three different protease digests. The aliquots were then digested separately overnight at 37°C with either Trypsin (Promega), Chymotrypsin (Promega), or Alpha lytic protease (New England Biolabs)) in a 1:30 (w/w) ratio. Desalting and peptide enrichment was performed using an Oasis HLB desalting 96-well µElution plate (Waters) attached to a vacuum manifold. The peptides were combined and analyzed by nanoLC-ESI MS with an Easy nLC 1200 (Thermo Fisher Scientific) system coupled to an Orbitrap Fusion mass spectrometer (Thermo Fisher Scientific) using stepped higher energy collision-induced dissociation (HCD). An EasySpray PepMap RSLC C18 column (75µm x 75 cm) was used to separate the peptides. A trapping column (PepMap 100 C18 3µM 75µM x 2cm) was used in line with the LC prior to separation with the analytical column. For LC separation, buffer A consisted of 0.1% formic acid and 80% acetonitrile in 0.1% formic acid. The LC conditions were as follows: 280-minute linear gradient consisting of 5-40% B in 0.1% formic acid over 240 minutes. The %B was then increased to 95% over 15 minutes and held for another 15 minutes before reducing the %B to 5%. The flow rate was set to 300 nL/min. The spray voltage was set to 2.5 kV and the temperature of the heated capillary was set to 55 °C. The ion transfer tube temperature was set to 275 °C. The scan range was 350−1800 m/z. Stepped HCD collision energy was set to 15, 25 and 45% and the MS2 for each energy was combined. Precursor and fragment detection were performed using an Orbitrap at a resolution MS1= 120,000. MS2= 30,000. The AGC target for MS1 was set to standard and injection time set to auto. Glycopeptide fragmentation data were extracted from the raw MS files using Byos (Version 5.5; Protein Metrics Inc). The glycopeptide fragmentation data were evaluated manually for each glycopeptide. The peptide was scored as true-positive when both the oxonium ions corresponding to the identified glycan and the correct b and γ fragment ions were observed. The Protein Metrics 305 N-glycan library with sulphated glycans added manually, was used to search the MS data. The relative amounts of each glycan at each site in addition to the unoccupied proportion was determined by comparing the extracted chromatographic areas for different glycoforms with an identical peptide sequence. A 1% False discovery rate (FDR) was applied, and the precursor mass tolerance was set at 4 ppm, and 10 ppm for fragments. All charge states for a single glycopeptide were summed. Glycans were categorized according to the composition detected. HexNAc(2)Hex(9-3) was classified as M9-M3, HexNAc(2)Hex(10+) was defined as M9GLc and any of these structures which contained fucose were characterised as FM (fucosylated mannose). HexNAc(3)Hex(5−6)X was classified as Hybrid with HexNAc(3)Hex(5-6)Fuc(1)X classified as Fhybrid. The complex-type glycans were categorised according to the number of HexNAc subunits and the presence or absence of fucosylation. Core glycans refer to truncated structures smaller than M3. M9glc-M4 were classified as oligomannose-type glycans. The oligomannose- and hybrid-type glycans were combined into a high mannose glycan category.

### Enzyme-Linked Immunosorbent Assay (ELISA)

ELISA assays were conducted using biotinylated proteins on streptavidin-coated plates, following previously established protocols. In brief, 96-well half-area clear plates (Corning, Thermo Fisher Scientific) were coated overnight at 4°C with 2 μg/mL streptavidin (Jackson ImmunoResearch, cat #016-000-113). The plates were then washed three times with 1X PBS/0.05% Tween (Sigma-Aldrich, cat #1003620819) and blocked with 100ul 3% BSA (Sigma-Aldrich, cat #A9418-500G) in PBS for 1 hour at room temperature (RT). BSA was dumped from the wells and plates were patted dry. Biotinylated proteins were added at a concentration of 2 μg/mL in 1% BSA/1X PBS/0.05% Tween and incubated for 1.5 hours at RT. Following incubation, plates were washed three times before adding diluted monoclonal antibodies (mAbs) or serum samples, which were incubated for an additional 1.5 hours. After washing, alkaline phosphatase-conjugated secondary antibodies (Jackson ImmunoResearch Laboratories, cat #109-055-170) were applied 50ul/well at 1:1000 dilution and incubated for 1 hour. Dissolve 1 tablet of phosphatase substrate (Thermo Fisher Scientific, cat #S0942-200TAB) per 5 ml of Alkaline staining buffer (1L recipe: add 2.03g MgCl2-6H2O (Fisher Bioreagents, cat #BP214-500) and 8.4g Na2CO3 (Sigma-Aldrich, cat #S7795-500G) in MilliQ water, adjust pH to 9.8, adjust volume to 1L with MilliQ water, add 1.0g NaN3 (Sigma-Aldrich, cat #S2002-100G), filter through 0.22μm filter, store at 4°C). Absorbance was measured at 405 nm using a Synergy HTX multi-mode reader after 20 minutes of substrate development.

### Cell Surface Binding Assay

HEK293T cells were seeded in 6-well plates and transfected with plasmids encoding the antigen of interest using Lipofectamine 2000 (Thermo Fisher Scientific, cat #11668500), following the manufacturer’s protocol. After 48 hours of incubation at 37°C with 5% CO₂, cells were harvested using FACS buffer (PBS + 2% FBS+5mM EDTA (Invitrogen, cat #15575-038), followed by washing twice with FACS buffer and resuspended at a density of 1 × 10L cells/mL. Cells were incubated with primary antibodies of interest at a final concentration of 10 µg/mL for 1 hour on ice. After incubation, cells were washed three times with FACS buffer and subsequently stained with Mouse Anti-Human IgG FC-PE (SouthernBiotech, cat #9040-09) for 1 hour on ice in the dark. Following a final set of washes, cells were resuspended in FACS buffer and acquired on a Bio-Rad ZE5 flow cytometer. Data were analyzed using FlowJo software.

### Biolayer Interferometry

For high throughput antigenicity screening of Q23-APEX immunogen designs and of isolated mAbs from immunized animals, BLI was performed with 10ug/ml IgG antibody. IgGs were immobilized on ProA sensors (Sartorius, cat #18-5012) to a signal of at least 1.0 nm using an Octet Red96 instrument (ForteBio). The immobilized IgGs were then dipped in the running buffer (1X PBS, 0.02% Tween20, pH 7.4) followed by 500 nM of trimers. Following a 120 s association period, the tips were dipped into the running buffer and dissociation was measured for 240 s. For assessing the polyclonal immune serum IgG responses in vaccinated animals, serum was used with a 1:10 dilution. Biotinylated trimers were first captured on SA biosensors (Sartorius, cat #18-5020) to a signal of at least 1.0 nm using an Octet Red96 instrument (ForteBio). The immobilized trimers were then dipped in the running buffer (1X PBS, 0.02% Tween20, pH 7.4) followed by polyclonal serum IgG. Following a 120 s association period, the tips were dipped into the running buffer and dissociation was measured for 240 s. For determining BLI kinetics with Fab versions of mAbs,, monoclonal IgG Fab heavy chain plasmids were engineered by inserting a His-Avi tag, followed by a stop codon, upstream of the disulfide bond in the Fc region. Paired heavy and light chain plasmids were co-transfected along with BirA for biotinylation into Expi293 cells (Thermo Fisher Scientific, cat #A14527) at a 2:2:1 ratio (HC:LC:BirA) using FectoPRO transfection reagent (Polyplus, cat #116-001). After 24 hours, the cells were supplemented with 0.3 M valproic acid (Sigma, cat #P4543-100G) and 40% glucose (Gibco, cat #A2494001). MAb IgG Fabs were harvested from the culture supernatant five days post-transfection by affinity chromatography using Ni Sepharose 6 Fast Flow (Cytiva, cat #17531802), according to the manufacturer’s instructions. The eluted antibodies were buffer exchanged into PBS and concentrated using a 10 kDa ultracentrifugal filter (Millipore, cat #UFC905024). Concentrated Fabs were subjected to size exclusion chromatography on a Superdex 200 Increase 10/300 GL column (Sigma-Aldrich, cat #GE28-9909-44). Specific fractions were pooled, further concentrated. BLI was performed with 10ug/ml Fab. Fabs were immobilized on SA sensors (Sartorius, cat #18-5020) to a signal of at least 1.0 nm using an Octet Red96 instrument (ForteBio). The immobilized Fabs were then dipped in the running buffer (1X PBS, 0.02% Tween20, pH 7.4) followed by 2-fold dilution of trimers starting at 500nM. Following a 120 s association period, the tips were dipped into the running buffer and dissociation was measured for 240 s.

### Neutralization Assay

Sera (1:10 starting dilution) or monoclonal antibodies (300 μg/ml starting concentration) were three-fold serially diluted in 25 µl of complete DMEM and incubated with HIV-1 Env-pseudotype virus (25 μl) for 60 minutes at 37°C in duplicate 96-well Culture Plates. TZM-bl cells (20,000 cells per well) with 20 μg/ml DEAE-Dextran were then added (50 μl) and incubated overnight. Control wells included cells only (background) and virus only (maximal entry). Serial dilutions were performed with tip changes to prevent carryover. After 72 hours, luciferase activity was measured using the Bright-Glo Luciferase Reporter Assay (Promega, cat #E2650) and a Synergy HTX multi-mode luminometer. Percent neutralization was calculated as: ((RLU_Virus_ – RLU_test_) / RLU_Virus_) × 100 Background RLU from uninfected control wells was subtracted before final calculations. Neutralizing serum titers (ID₅₀) and antibody titers (IC₅₀) were determined via a four-parameter nonlinear dose-response inhibition curve.

### PCT64^LMCA^ knock-in mouse adoptive transfer immunization study

Previously described PCT64^LMCA^ knock-in (KI) mice ^42^ donor derived CD45.2 B cells were isolated from spleens using Pan B Cell isolation kit (Milteny Biotec) following manufacturer’s protocol. Cells were counted using Luna FL cell counter. B cells were resuspended in PBS and 1 × 10^5^ B cells were injected intravenously in tail vein of recipient 8-12 weeks old B6.SJL-Ptprca Pepcb/BoyJ (CD45.1 mice). 1 day post adoptive transfer groups of recipient animals (n = 5 animals per group) were immunized either with SMNP adjuvant only (control 5 μg) or 20 μg of Q23-APEX-GT2 trimer protein adjuvanted with 5 μg of SMNP subcutaneously (SC) at the base of the tail or 1μg of Q23-APEX-GT2 mRNA lipid nanoparticle (LNPs) (provided by Moderna) intramuscularly (IM) in each hindleg. For animals receiving SC immunization draining inguinal lymph nodes and for animals receiving IM immunization iliac and popliteal lymph nodes were harvested at week 2, 4 post immunization. Immune sera at pre-immunization, weeks 2 and 4 were also collected.

### Analysis of PCT64^LMCA^ knock-in mouse B cell responses

Single cell suspension was prepared by gently crushing lymph nodes and passing them through a 70 μM strainer. Incubation with PBS containing Live/Dead Blue (Thermo Scientific) diluted 500-fold and FcR Blocking reagent (Purified Rat anti-mouse CD16/CD32, BD Biosciences) diluted 200-fold was done for 20 mins at 4°C. After washing, BCR antigen staining was done using biotinylated Q23-APEX-GT2 trimer conjugated to either streptavidin-BV510 (BioLegend) or strepatvidin-Alexa647 (BioLegend) for 30 mins at 4°C. Excess antigen was washed off and, cells surface staining was done with an antibody cocktail containing CD4, CD8, F4/80, GR-1, NK1.1 (APC-eFluor 780, eBioscience, clone RM4-5, 53-6.7, BM8, RB6-8C5, PK136 respectively), B220 (BUV395, BD Bioscience, clone RA3-6B2), CD38 (BUV563, BD Biosciences, clone 90), CD95 (PE-Cy7, BioLegend, clone L138D7), CD45.1 (BV605, BioLegend, clone A20), CD45.2 (PE, BD Biosciences, clone 104), CD138 (BV650, BD Biosciences, clone 281-2), IgD (Alexa 594, Biolegend, 11-26c.2a clone) and IgM (BV750, II/41 clone) for 30 mins at 4°C. Flow cytometry data was acquired using BD FACS Symphony A5 cell analyzer. For cell sorting, Live/Dead stain was replaced with SYTOX Green (Thermo Fisher Scientific). The antibodies used for sorting were CD4, CD8, F4/80, GR-1 & NK1.1 (APC-eFluor 780, eBioscience, clone RM4-5, 53-6.7, BM8, RB6-8C5, PK136 respectively), B220 (Alex Fluor594 BioLegend, clone RA3-6B2), CD38 (BB700, BD Biosciences, clone 90), CD95 (PE-Cy7, BioLegend, clone L138D7), CD45.1 (APC R700, BD Biosciences, clone A20), CD45.2 (PE, BD Biosciences, clone 104), IgD (BV605, BioLegend, clone 11-26c.2a), CD138 (BV650, BD Biosciences, clone 281-2). Cells from each individual mouse were barcoded with TotalSeq™-C anti-mouse Hashtag Antibodies. A total of 10 hashtags were used. Cells were washed 3 times to remove any excess antibodies.

### Cell sorting and paired BCR sequencing of PCT64^LMCA^ B cell responses

Cells were sorted using BD FACS symphony S6 using 85μM nozzle. Samples were sorted onto PCR tubes with PBS buffer containing 10% FBS. Encapsulation of sorted cells and NGS library preparation was performed following the 10x Genomics Chromium Next GEM Single Cell 5’ Reagent Kits v2 protocol (10x Genomics). TapeStation Systems D5000 high sensitivity Screen Tape assay (Agilent, Santa 5 Clara, CA) was used to measure library size. After quantifying the libraries through Qubit dsDNA High Sensitivity (Invitrogen), they were pooled and were run on NextSeq 550 System (Illumina, San Diego, CA). Analysis was performed using Cell Ranger v.6 software pipeline (10x Genomics) with a customized reference database. Sequencing data was analyzed using Geneious Prime software (Geneious) and IMGT/V-Quest.

### Immunization in rhesus macaques, blood and lymph node processing

2 groups of 6 rhesus macaques each, evenly distributed by gender between the ages 3-5 years, were immunized with germline-targeting trimer, Q23-APEX-GT2 trimer protein + SMNP adjuvant^64^ (Arm 1) and membrane-anchored mRNA LNPs (Arm 2). Rhesus macaques were primed at week 0 with 100 μg Q23-APEX-GT2 protein + SMNP adjuvant (Arm 1) or 100 μg Q23-APEX-GT2 mRNA LNPs (Arm 2) administered subcutaneously and distributed into four injection sites (split equally between bilat inner mid-upper arm and bilat inner mid-thighs, 25 μg each). The Arm 1 Q23-APEX-GT2 protein priming immunization was given as escalating dose (DE) over 2 weeks ^65,66^ and a single bolus dose for Q23-APEX-GT2 mRNA LNPs in Arm 2. For escalating dose priming, animals received seven injections of the Q23-APEX-GT2 protein along with the SMNP adjuvant across four sites over 12 days (on days 0, 2, 4, 6, 8, 10, and 12). The total Q23-APEX-GT2 trimer immunogen doses at each injection were: 0.2, 0.43, 1.16, 3.15, 8.56, 23.3, and 63.2 µg, evenly distributed across the four immunization sites. Protein prime was co-administered with 375 µg of SMNP adjuvant, which was delivered in a proportional dose-escalation manner alongside the seven protein immunogen injections. Both groups were bolus boosted with 100 μg of Q23-APEX-GT2 protein + SMNP adjuvant (Arm 1) or Q23-APEX-GT2 mRNA LNPs (Arm 2).

Peripheral blood was collected in sterile vacutainers containing acid citrate dextrose formula A (ACD-A) as an anticoagulant (DB Vacutainer cat #364606). A total of 40 mL of ACD-A blood was centrifuged at 1000g for 10 minutes at 20°C in sterile 50 mL conical tubes. Plasma was collected without disturbing the buffy coat or red blood cell pellet, then subjected to a second centrifugation at 1500g for 15 minutes at 20°C to remove all cellular material. The cell-depleted plasma was aliquoted into 1 mL cryovials (Sarstedt cat # 72.694.396) and stored at −80°C. The cell fraction was resuspended in an equal volume of Hanks’ Balanced Salt Solution (HBSS) without calcium or magnesium (HBSS−/−) (Gibco cat # 14175-079) containing 2 mM EDTA (Invitrogen cat #15575-020) and divided into four 50 mL conical tubes. Additional HBSS−/− with EDTA was added to each tube to bring the total volume to 35 mL. The cell suspension was under layered with 14 mL of 96% Ficoll-Paque Plus (Cytiva cat # 17144003) and centrifuged at 725g for 20 minutes at 20°C with slow acceleration and braking. Mononuclear cells at the Ficoll interface were collected, transferred to a fresh 50 mL conical tube containing HBSS−/− with EDTA, and washed by centrifugation at 200g for 15 minutes at 20°C. Following removal of the supernatant, the cell pellet was resuspended in 40 mL of HBSS containing calcium and magnesium (HBSS+/+) (Gibco cat # 24020-117) supplemented with 1% fetal bovine serum (FBS) (Cytiva cat # SH300.71.03). The suspension was centrifuged at 200g for 15 minutes at 20°C, after which the supernatant was discarded. This centrifugation step effectively pelleted white blood cells (WBCs) while leaving most platelets in suspension. The mononuclear cell pellet was gently resuspended in the residual media, followed by the addition of HBSS+/+ with 1% FBS to a final volume of 10 mL. Cells were counted, and viability was assessed using ViaStain AOPI solution (Revvity cat #CS2-0106-25ml) and a Cellometer Auto 2000 instrument (Revvity, Waltham MA). The cells were then centrifuged at 300g for 10 minutes at 20°C, the supernatant was discarded, and the pellet was resuspended in CryoStor CS5 cryopreservation medium (Stemcell technologies cat # 07930) at a final concentration of 5–10 × 10L cells/mL. The suspension was aliquoted into 1 mL cryovials (Thermoscientific cat # 374503), stored in a Corning CoolCell LX (Corning cat #432002) or FTS30 (Corning cat #432006) freezing container at −80°C overnight, and subsequently transferred to vapor-phase liquid nitrogen for long-term storage. Mononuclear cells from lymph nodes (LNs) were processed in a similar protocol to blood mononuclear cells. Axillary and/or inguinal LNs were excised, placed immediately into RPMI1640 medium (Corning cat # MT15040CV) on wet ice and processed within 6 hours. LNs were diced with a sterile scalpel and passed through a sterile cell strainer (Falcon cat # 352360). Cells were collected from the pass-through and subjected to Ficoll density gradient purification as described above, with RPMI1640 supplemented with 10% FBS used instead of HBSS.

### Flow cytometry antigen-specific B cell sorting of macaque lymph node samples

Avi-tagged, biotinylated Q23-APEX-GT2 trimer and its double knockout variant (169E-171E) were conjugated to streptavidin-labeled fluorophores at room temperature for 30 minutes. Cryopreserved LN samples were thawed and resuspended in RPMI medium (Thermo Fisher, cat # MT15040CV) supplemented with 50% FBS. Cells were washed with FACS buffer (PBS + 2% FBS + 2mM EDTA) (Invitrogen, cat #15575-038) and stained with fluorescently labeled anti-CD3 (BD Pharmingen 557757), CD4 (Biolegend 317418), CD8 (Biolegend 557760), CD14(BD Pharmingen 561384), CD19 (Biolegend 302230), CD20 (Biolegend 302326), IgG(BD Horizon 564230, and IgM(Biolegend 314508) antibodies for 30 minutes on ice in the dark. The conjugated antigens were then added and incubated for another 30 minutes. A 1:250 dilution of FVS510 LIVE/DEAD stain (Thermo Fisher Scientific, cat #L34966) was applied, followed by a 15-minute incubation. Before sorting, cells were washed with FACS buffer and passed through a cell strainer into a 5 mL round-bottom tube (Corning 352058). Sorting was conducted on a BD FACSMelody, and cells were collected into 96-well PCR plates for single-cell sequencing. RNA extraction, cDNA synthesis, and VDJ gene amplification were performed as previously described. Briefly, RT-PCR was performed using Superscript IV reaction and IgH, IgK, and IgL primers. Paired HC and LC sequences were amplified using nested PCR reactions and analyzed by 2% 96 E-gels with SYBR Safe (Thermo Fisher Scientific cat #G720802). Wells with successful DNA PCR amplification were Sanger sequenced (Azenta).

### Antibody Cloning

IgG heavy and light chain sequences for sorted B cell derived antibodies with features characteristic of rhesus V2-apex bnAbs from all immunized animals were cloned in IgG1, IgK, IgL AbVec Vector upstream of respective constant region using AgeI/NheI, AgeI/BsiWI and AgeI/SaII restriction sites respectively.

### Monoclonal Antibody Expression and Purification

Paired heavy and light chain plasmids for Rhesus and human V2-apex UCAs or monoclonal antibodies cloned from immunized animals were co-transfected in Expi293 cells (Thermo Fisher Scientific, cat #A14527) in a 1:1 ratio using FectoPRO (Polyplus, cat #116-001) transfection reagent and were fed with 0.3M valproic acid (Sigma, cat #P4543-100G) and 40% Glucose (Gibco, #A2494001) 24 hours after the transfection. Monoclonal IgGs were purified from the culture supernatant five days post-transfection with 1:1 ratio Protein-A (Cytiva, cat #17127903) and Protein-G Sepharose beads (Cytiva, cat #17061805) per manufacturer’s instructions. After elution with IgG elution buffer (Thermo Fisher Scientific, cat #PI21009), antibodies were buffer exchanged into PBS using 50kDa Ultra centrifugal filter unit. (Millipore, cat #UFC905024)

### Bulk Repertoire Sequencing and Immunogenetic Analysis

Cryopreserved lymph node samples were rapidly thawed in a 37°C water bath for 1–2 minutes until a small ice pellet remained. The thawed cells were immediately transferred to pre-warmed RPMI medium (Thermo Fisher, cat #MT15040CV)) supplemented with 50% FBS in a dropwise manner while gently swirling to minimize osmotic shock. Cells were centrifuged at 400 × g for 5 minutes at room temperature, and the supernatant was carefully aspirated to remove residual cryoprotectants. The cell pellet was resuspended in fresh RPMI medium, and viability was assessed using trypan blue exclusion (Sigma T8154). For RNA extraction, cells were lysed in RLT Plus buffer (Qiagen, cat #1053393) supplemented with β-mercaptoethanol, and total RNA was extracted using the RNeasy Plus Mini Kit (Qiagen, cat #74134) according to the manufacturer’s instructions, including on-column DNase I treatment (Qiagen cat #79254) to eliminate genomic DNA contamination. RNA quality and concentration were assessed using a NanoDrop spectrophotometer (Thermo Fisher Scientific) and the Agilent 2100 Bioanalyzer RNA Integrity Number (RIN) score. Only samples with RIN ≥ 7 were used for downstream applications. Reverse transcription was performed using gene-specific primers targeting IgM and IgG constant regions. cDNA synthesis was carried out with SuperScript IV reverse transcriptase (Thermo Fisher Scientific, cat #1750150) following the manufacturer’s protocol, with an initial primer annealing step at 65°C for 5 minutes, followed by reverse transcription at 55°C for 60 minutes and enzyme inactivation at 70°C for 15 minutes. The resulting cDNA was purified using ExoSAP-IT (Thermo Fisher Scientific, cat #78205) to remove excess primers and dNTPs. Immunoglobulin heavy-chain variable region (VDJ) amplification was performed in two sequential PCR steps using HotStarTaq Plus DNA Polymerase (Qiagen, cat #203603). The first PCR utilized a set of framework region primers spanning VDJ segments, followed by a nested PCR with primers incorporating Illumina-compatible overhang sequences. PCR products were enzymatically cleaned using ExoSAP-IT, and Illumina sequencing adapters with unique dual indexes were introduced via a second round of PCR. Final DNA libraries were purified using SPRIselect beads (Beckman Coulter Genomics, cat #B23318) with a 0.8x bead-to-sample ratio to remove adapter dimers and short fragments. Library concentration was quantified using a Qubit fluorometer (Thermo Fisher Scientific), and fragment size distribution was assessed using a Bioanalyzer (Agilent 2100) High Sensitivity DNA chip. Sequencing libraries were pooled in equimolar concentrations and loaded onto an Illumina NovaSeq 6000/NextSeq 1000/2000 system using a 2 × 300 bp paired end read configuration.

### B-cell Annotation / VDJ Repertoire Analysis

Filtered and quality-controlled IgM and IgL sequences were analyzed using IgDiscover v0.15.1 to curate a personalized immunoglobulin repertoire library for each rhesus macaque (Table S3). Merged reads served as input, with the KIMDB 1.1 database (http://kimdb.gkhlab.se/) as the reference for heavy chains and the Ramesh et al. allele database for light chains. To enhance accuracy, the IgDiscover J output for both heavy and light chains was further processed using the “discoverjd” feature, applying a J coverage threshold of 100 and an allele ratio cutoff of 0.33 to exclude low-confidence novel alleles. This approach enabled the identification of both known and novel V and J gene alleles for each rhesus macaque, confirmed the presence of known D gene alleles, and facilitated the precise assignment of V, D, and J alleles for each broadly neutralizing antibody lineage.

### Antibody Clonotypes

Antibody clonotypes were defined as sequences sharing the same V and J germline segments and identical CDRH3 amino acid sequences to minimize sequencing and amplification errors. While collapsing V and J regions to germline assignments prevents double-counting errors in these regions, it does not address errors in CDRH3. To assess their impact on clonotype diversity, we grouped sequences into clonotypes with no mismatches in the CDRH3 sequence.

### Ployclonal Fab preparation

To generate polyclonal serum Fabs for EMPEM studies, poly-IgG was first isolated from immune sera by rotating 1ml of serum with 0.25ml of Protein A Affinity Chromatography Resin (Cytiva, cat #17127903) and 0.25 ml Protein G Sepharose (Cytiva, cat #17061805), and 9ml PBS. The next day, beads were transferred to an Econo-Pac column (Bio-Rad Laboratories), washed with 2 column volumes of PBS, and then eluted with 10ml IgG elution buffer (Thermo Fisher Scientific, cat #PI21009). After elution, using 50kDa Ultra Centrifugal Filter Units (Merck Millipore cat #UFC9030) to buffer exchange poly-IgG Abs into PBS and then concentrated. Polyclonal antibodies and mAbs were then digested using Pierce™ Fab Preparation Kit (Thermo Fisher cat #44985) following manufacturer instructions. After digestion, 10 kDa Ultra Centrifugal Filter Units (Millipore, cat# UFC9010) were used to buffer exchange proteins into PBS and concentrate to 0.5ml. Concentrated Fabs underwent size exclusion chromatography using Superdex 200 Increase 10/300 GL column (Sigma-Aldrich, cat #GE28-9909-44). Specific fractions from the chromatography run were mixed and concentrated before being used.

### EMPEM data collection and processing

15 µg of Q23-APEX-GT2 trimer was incubated with 500 µg of polyclonal Fab overnight at room temperature before purifying through size exclusion chromatography with a Superdex 200 Increase 10/300 column (GE Healthcare). The complex was diluted to 0.02 mg/ml in 1x Tris-buffered saline pH 7.4, and 3 µl were applied to a 400 mesh Cu grid, blotted with filter paper and stained with 2% uranyl formate or NanoW (Nanoprobes). Samples were imaged on either a 200 kV Thermo Fisher Scientific Glacios with a Falcon IV direct electron detector (1.89 Å/pixel; 73,000x magnification) or a 200 kV Thermo Scientific Talos F200C with a Ceta 16M camera (2 Å/pixel; 73,000x magnification) using EPU software. Particle picking, 2D classification and 3D reconstructions were done using Relion 4.0^90^. Following three rounds of 2D classification, particles from classes corresponding to immune complexes were subjected to 3D refinement with C3 symmetry and a 40 Å low-pass filtered map of HIV Env ectodomain as the initial model. The initial model is based on PDB coordinates 6V0R, converted in a map using the molmap feature in UCSF ChimeraX^91^. Following 3D refinement, C3 symmetry expansion was applied to the particles and 7 separate focused 3D classification “skip align” jobs were run (K=10), each with a 40 Å diameter spherical mask over key HIV Env epitopes. The names of the epitopes and reference structures used for orienting the masks are: 1) gp41-base (PDB 6X9R), 2) gp41-GH (PDB 7L8U), 3) gp41-FP (PDB 7L8T), 4) gp120-GH (PDB 7L8B), 5) C3V5 (PDB 7L86), 6) CD4bs and gp120 interface (PDB 7L8X), and 7) V1V2V3 (PDB 7L8E). For each epitope 3D classification, classes with visible Fab density were selected and subjected to 3D refinement, 2D classification, and a second round of 3D classification. If the 3D refinement resulted in partial Fab density relative to the Env trimer, classes were selected from the subsequent round of 3D classification, and this was repeated until the reconstruction improved, or no change was noted. Final reconstructions were visually inspected and assigned the correct epitope label. The number of final particles belonging to each epitope was divided by the total number of particles in the initial (C3) 3D refinement. This value, which can range (per epitope) from 0 to 3 due to C3 symmetry expanded particles used in the 3D classification steps, is described as the *EMPEM magnitude*. The sum of each individual *EMPEM magnitude* is the *Total EMPEM magnitude*. 3D maps were segmented and figures generated using UCSF Chimera^92^. Fab densities that could not be resolved in 3D are presented as false-colored 2D classes. Representative EM maps have been deposited to the Electron Microscopy Data Bank (EMDB).

### Monoclonal Fab preparation for Cryo-EM structural studies

Monoclonal IgG Fab heavy chain plasmid was designed by inserting a His-Avi tag followed by a stop codon before the disulfide bond in the Fc region. Paired truncated heavy and light chain plasmids were co-transfected in Expi293 cells (Thermo Fisher Scientific, cat #A14527) in a 1:1 ratio using FectoPRO (Polyplus, cat #116-001) transfection reagent and were fed with 0.3M valproic acid (Sigma, cat #P4543-100G) and 40% Glucose (Gibco, #A2494001) 24 hours after the transfection. Fabs were purified from the culture supernatant five days post-transfection with Ni Sepharose 6 Fast Flow (cytiva, cat#17531802) per manufacturer’s instructions. After elution, Fab proteins were buffer exchanged into PBS and concentrated using 10kDa Ultra centrifugal filter unit. (Millipore, cat #UFC905024) Concentrated Fabs underwent size exclusion chromatography using Superdex 200 Increase 10/300 GL column (Sigma-Aldrich, cat #GE28-9909-44). Specific fractions from the chromatography run were pooled and concentrated before being used.

### Cryo-EM data collection and processing

The high-resolution unbound (apo) and Fab-bound structures of Q23-APEX-GT2 envelope trimer were determined using single-particle cryo-EM as previously described^93^. Samples were prepared by diluting purified Q23-APEX-GT2 to 2.5 mg/mL with PBS or 3-fold molar excess of Fab:trimer and incubated on ice for 30 minutes, and then supplemented with a final concentration of 0.005% (w/v) n-Dodecyl β-D-maltoside (DDM) to prevent preferred orientation. Copper C-flat Holey carbon-coated grids (CF-1.2/1.3 300 mesh; EMS) were first glow discharged and then applied with 3 uL of sample for 30 seconds in a Vitrobot Mark IV at room temperature with 100% humidity. Sample grids were then vitrified using liquid ethane. Cryo-EM data were collected on a FEI Titan Krios 300 kV cryo-transmission electron microscope equipped with a Gatan K3 detector operating in counting mode using Leginon^94^. Defocus values were set to cycle between −0.80 and −2.0 μm, and a total dose of 58 e^-^/Å^2^ was fractionated over 50 raw frames. All processing was done in cryoSPARC v3.4^95^, including motion correction, CTF estimation, non-templated blob particle picking, 2D classification, *ab initio* modeling, and iterative 3D refinements. All homogenous and non-uniform 3D refinements of Fab-bound complexes were performed using C1 symmetry, while all homogenous 3D refinements and non-uniform 3D refinements for apo Q23-APEX-GT2 were performed with C1 and C3 symmetry, respectively. Data acquisition and processing details for all structures are provided in Dataset S1.

### Atomic model building and refinement

Each atomic model was solved by iterative manual rebuilding in Coot^96^ and real-space refinement in Phenix ^97^, with the overall structure quality for all models periodically assessed using MolProbity ^98^ and EMRinger ^99^ during refinement until satisfactory validation of each model was achieved. The initial model used for apo Q23-APEX-GT2 was another Q23.17-based prefusion-stabilized envelope trimer (PDB-7LLK) ^100^, which was fit into our 2.9 Å cryo-EM reconstruction density using UCSF ChimeraX ^101^. For atomic building of Fab complexes, the initial model of each Fab variable region (Fv) was first obtained using both the AlphaFold3 server ^102^ and the AbodyBuilder2 application of the SAbPred Antibody Prediction Toolbox ^103^; each Fv model for a specific Fab was fit into the respective cryo-EM 3D reconstruction using ChimeraX and the initial model bearing the HCDR3 with the best fit HCDR3 was selected for model building along with our apo Q23-APEX-GT2 structure. Protein interface calculations were performed using PDBePISA ^104^. Final model statistics and validations are provided in Dataset S1.

### Statistical Analysis

Statistical analyses were performed in GraphPad Prism and R. Statistical significance was assessed using a two-tailed Student’s t-test. P < 0.05 was considered statistically significant. *P < 0.05; **P < 0.01; ***P < 0.001; ****P < 0.0001; ns, no significant difference.

## Data availability

The cryo-EM structures of the HIV trimer alone or in complex with mAbs presented in this manuscript can be found in the Electron Microscopy Data Bank or Protein under accession codes; EMDB: EMD-49865 (PDB: 9NVV), EMD-49866 (PDB: 9NVW), EMD-49867 (PDB: 9NVX), EMD-49868 (PDB:9NVY), EMD-49869 (PDB: 9NVZ), EMD-49870 (PDB: 9NW0), EMD-49871 (PDB: 9NW1). The EMPEM structures can be found in the Electron Microscopy Data Bank under accession codes EMD-49511, EMD-49512 and EMD-49513. All sequencing data generated in this study have been deposited in the NCBI database under BioProject PRJNA1229425, with BioSample accessions SAMN47126428-SAMN47126475 and associated SRA files (SRX27822716-SRX27822762). The BCR sequencing data from Q23-APEX-GT2-immunized PCT64 LMCA mice are available under GenBank accessions PV237254-PV239402, while the sorted BCR sequences from Q23-APEX-GT2-immunized Rhesus Macaques are available under accessions PV256748-PV257578. All code used for data analysis in this study is publicly accessible on GitHub at https://github.com/nmishra93/gt2_rhesus_immunization.

## SUPPLEMENTARY FIGURES AND LEGENDS

**Figure S1.**
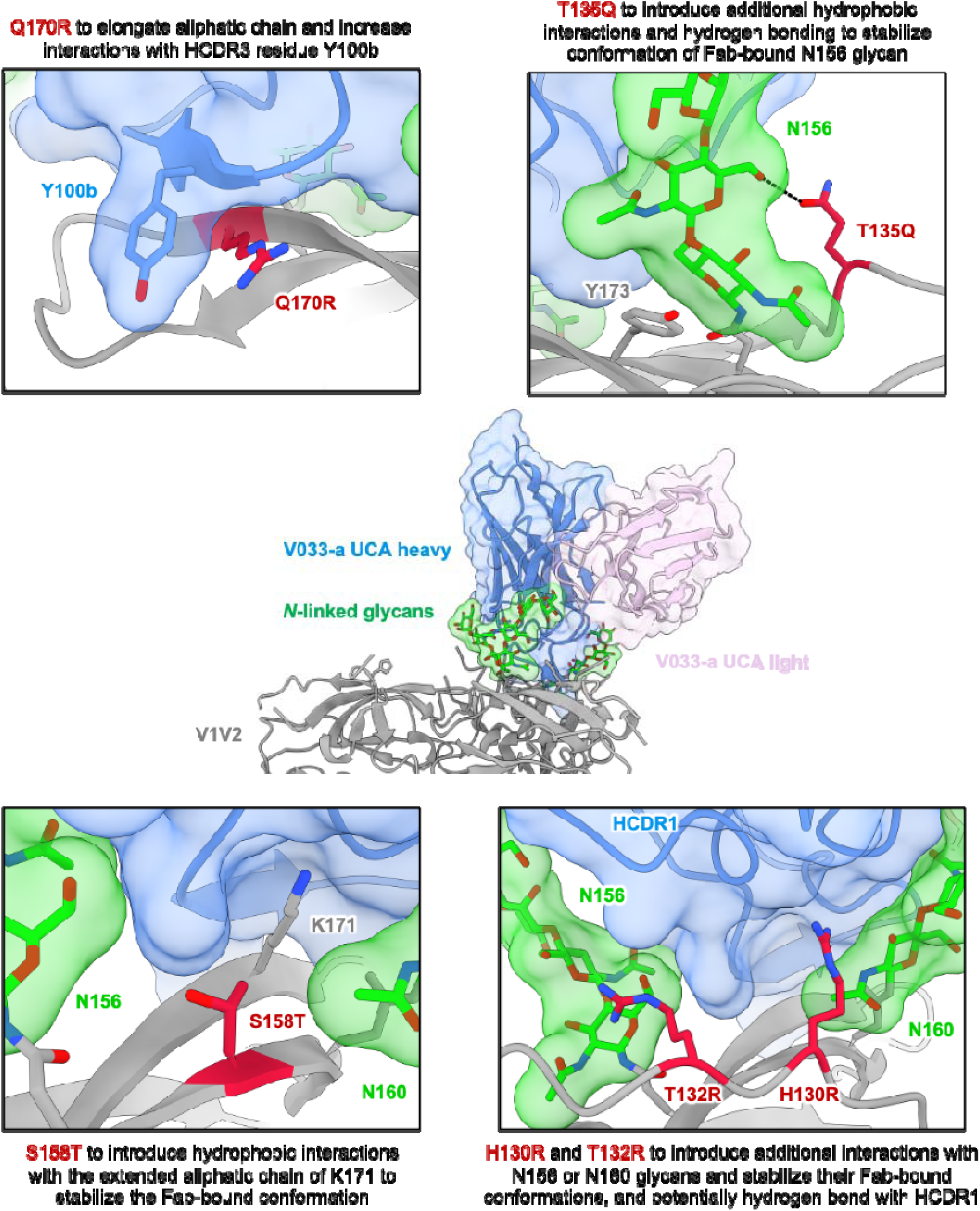
Antibody-guided design of V2-apex germline-targeting modifications. Related to Figure 1. The cryo-EM structure of the rhesus bnAb lineage V033-a unmutated common ancestor (UCA) in complex with Q23.17 MD39 envelope trimer was used to rationally design V2-apex germline-targeting modifications (PDB-XXXX) (Habib et al, 2025 *in preparation*). Expanded view in middle diagram highlights V2-apex binding orientation of the V033-a UCA Fab and contacted apical glycans, both of which are shown with transparent surface representation and colored according to labels (heavy chain, blue; light chain, pink; glycans, green). Insets depict the local environments of select mutations, which are shown in stick representation and colored red, to highlight examples of germline-targeting modifications engineered to increase the affinity of envelope for the V033-a UCA.

**Figure S2.**
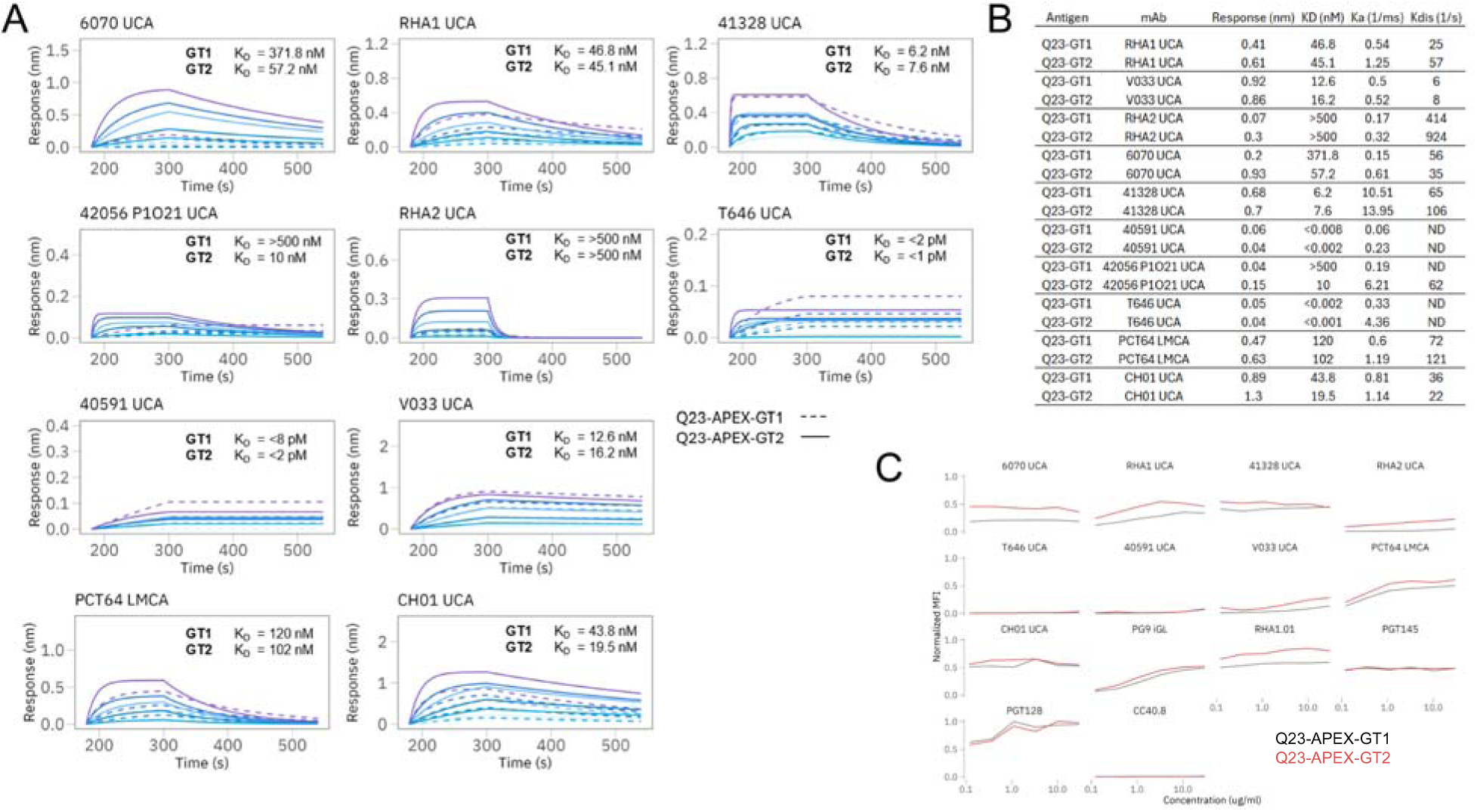
Engineered Q23-APEX-GT2 shows broad and enhanced binding to unmutated common ancestor versions of rhesus and human V2-apex bnAb. Related to Figure 1. A. Bio-Layer Interferometry (BLI) binding curves for Q23-APEX-GT1 and Q23-APEX-GT2 trimers against selected V2-apex unmutated common ancestors (UCAs) as Fabs. Fabs were tested at five two-fold serial dilutions, starting from a maximum concentration of 10 µg/ml. The highest concentration (10 µg/ml) is depicted in violet, while the lowest concentration (0.625 µg/ml) is shown in light blue. Binding kinetics were assessed to determine affinity and potential differences in epitope engagement between the engineered Q23-APEX-GT2 and the base construct, Q23-APEX-GT1 germline-targeting (GT) immunogens. BLI binding sensograms traces depict association and dissociation phases, with relative binding strength inferred from response units over time. B. K_D_, K_on,_ and K_off_ values with the maximum binding response (nm) values are shown as a table for all 10 UCAs with Q23-APEX-GT1 and Q23-APEX-GT2 trimers (from panel A). C. Antigenic assessment of membrane-bound Q23-APEX-GT1 and Q23-APEX-GT2 was conducted using a cell surface binding assay. The trimer immunogens were expressed on the cell surface via the wild-type (WT) Q23 transmembrane domain, mimicking their native presentation on viral membranes. Antibody binding was measured across a dilution series, starting at 10 µg/ml, to compare binding efficiencies and potential epitope accessibility in the transmembrane bound trimer context.

**Figure S3.**
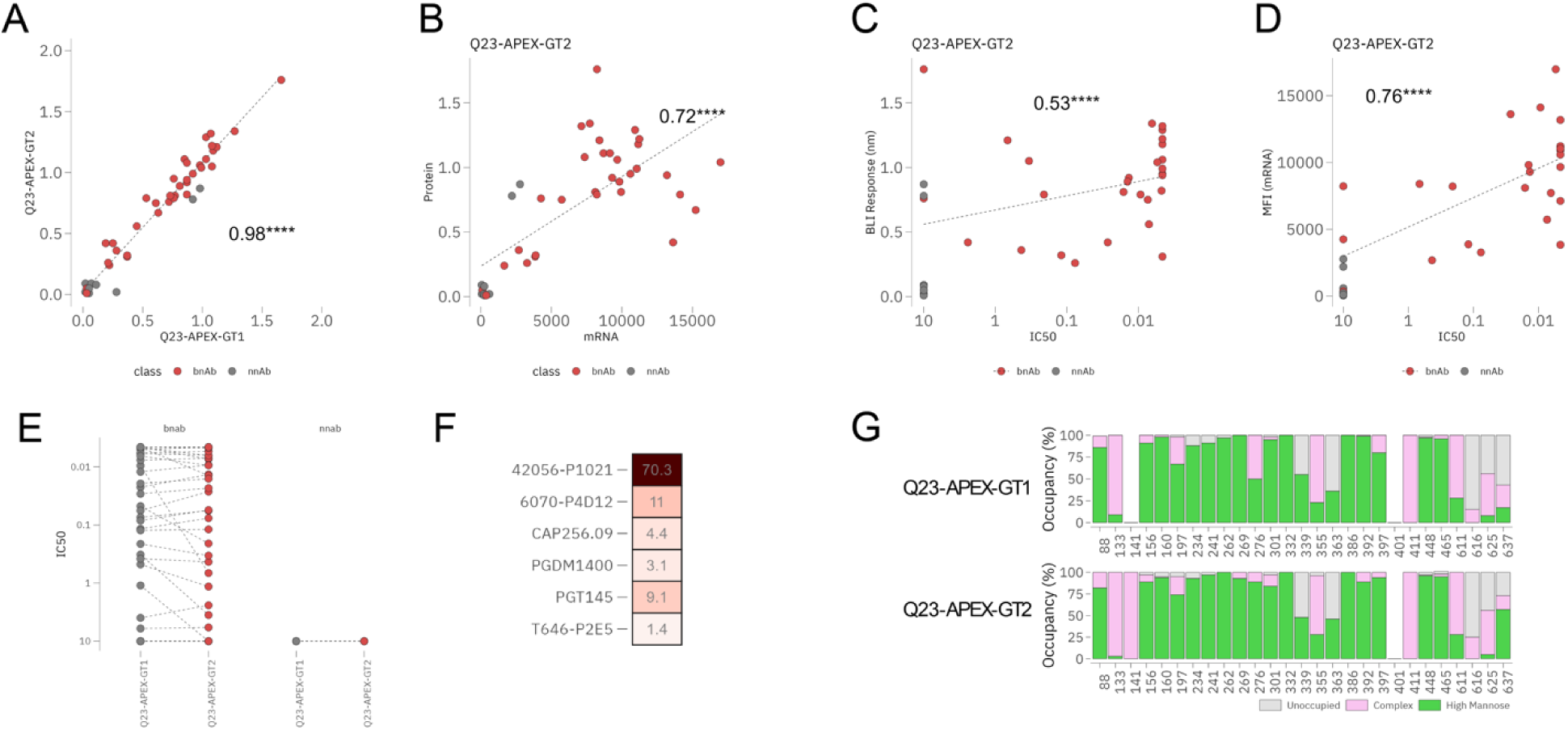
Engineered Q23-APEX-GT2 trimer shows native-like antigenicity and glycan profiles favorable for immunogenicity evaluation studies. Related to Figure 1. A. Antigenic characterization of Q23-APEX-GT2 and Q23-APEX-GT1 trimers in their soluble protein forms was evaluated using biolayer interferometry (BLI) against a comprehensive panel of broadly neutralizing antibodies (bnAbs, n = 36) and non-neutralizing antibodies (nnAbs, n = 11). The binding profiles indicate that both immunogens exhibit overall similar antigenic properties (R^2^ = 0.98, P <0.0001), suggesting retention of key epitope structures. B. Antigenic characterization of Q23-APEX-GT2 in its soluble protein form versus its membrane-bound presentation via mRNA expression. The soluble protein was analyzed by BLI, while the mRNA-expressed antigen was evaluated using a cell surface binding assay. Antibody binding readouts at 10 µg/ml were used for direct comparison between the two formats, highlighting comparable antigen exposure and accessibility when presented as a recombinant protein versus a cell-surface expressed antigen (R^2^ = 0.72, P <0.0001). C. Correlation plots for neutralization (IC_50_) versus BLI binding to soluble antigen, and (D). IC_50_ versus binding to cell surface expressed antigen shows a significant positive correlation (Pearson’s coefficient > 0.5). Four asterisks denote P value less than 0.0001. E. Q23-APEX-GT2 showed a reduction in IC_50_ (suggesting a slightly more resistant neutralization profile) for a few bnAbs targeting the V2-apex and CD4bs. (F). Fold reduction in IC_50_ values for several V2-apex bnAbs (from both human and rhesus) was observed for Q23-APEX-GT2 pseudoviruses compared to Q23-APEX-GT1, suggesting the acquisition of a difficult-to-neutralize phenotype. Neutralization assays were performed using a large panel of 70 monoclonals spanning several bnAbs and non-nAbs (starting at 10 µg/ml) targeting all major epitopes against both Q23-APEX-GT1 and Q23-APEX-GT2 pseudoviruses. G. Glycan profile characterization of Q23-APEX-GT1 and Q23-APEX-GT2, assessed through proteomics-based site-specific glycan analysis (SSGA). The composition of glycans at key glycosylation sites was analyzed, highlighting variations in glycan processing at the V2-apex glycans between the two constructs. High-mannose glycans are represented in faded green, complex-type glycans in light pink, and unoccupied glycosylation sites in gray.

**Figure S4.**
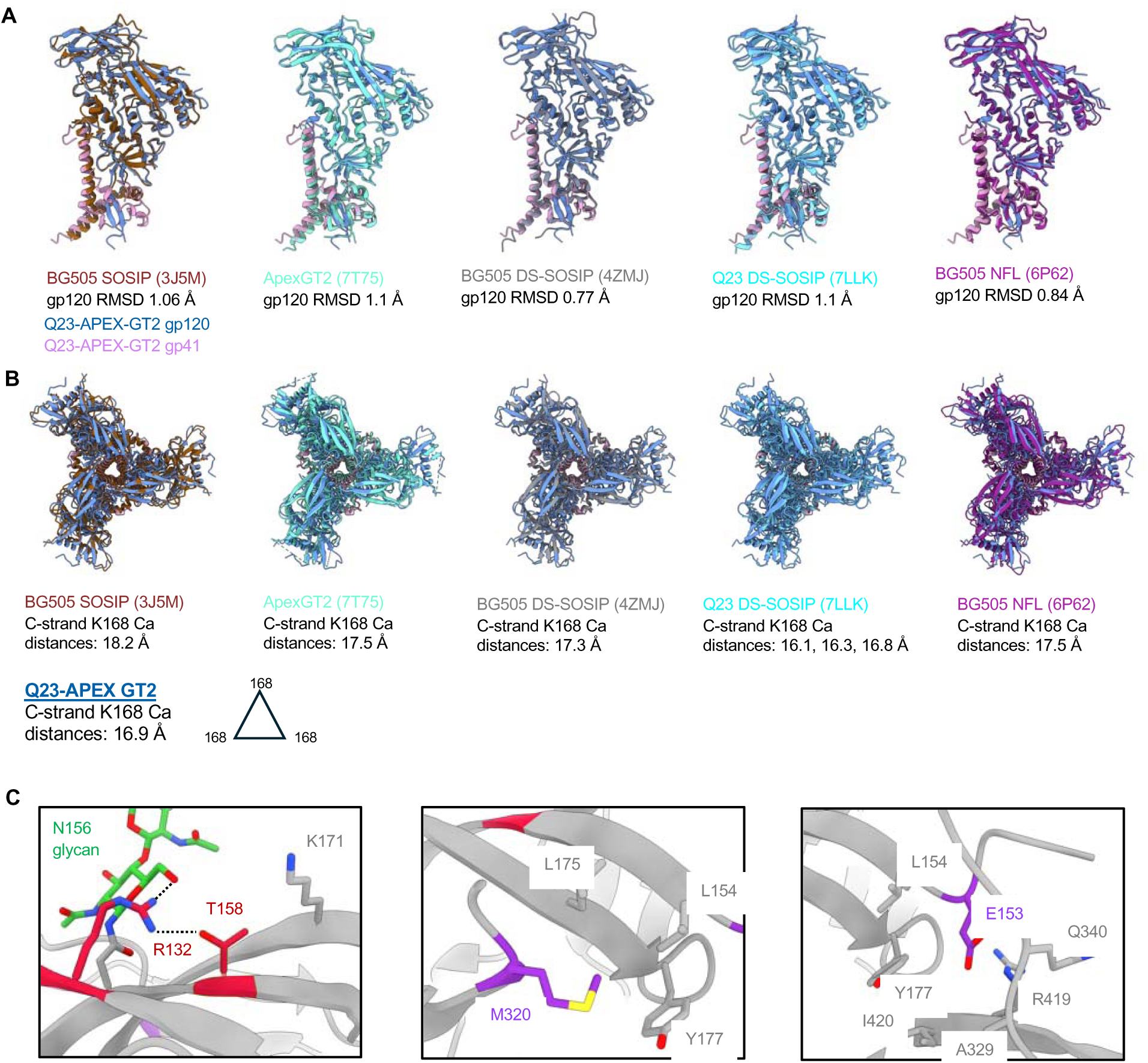
Structural alignments and details for Q23-APEX-GT2’s V2-apex bnAb site. Related to Figure 1. A. Superimpositions of a single protomer from Q23-APEX-GT2 with a protomer from the indicated envelope trimer structures by gp120 alignment. B. Superimpositions of Q23-APEX-GT2 with the indicated envelope trimers achieved by gp120 alignment. Prefusion-closed trimer distances are measured from the alpha carbon (Ca) of K168 residues between adjacent C-strands. C. Local environments of select GT1 (purple) and both GT2 (red) modifications in Q23-APEX-GT2. T132R forms hydrogen bonds with N156 glycan and S158T. The extended aliphatic chain of T132R makes hydrophobic interactions with N156, while the methyl group of S158T makes hydrophobic interactions with K171. T320M fills a hydrophobic pocket lined by V2 residues L154, L175, and Y177. G153E inserts into a large pocket formed by L154, Y177, A329, Q340, R419, and I420.

**Figure S5.**
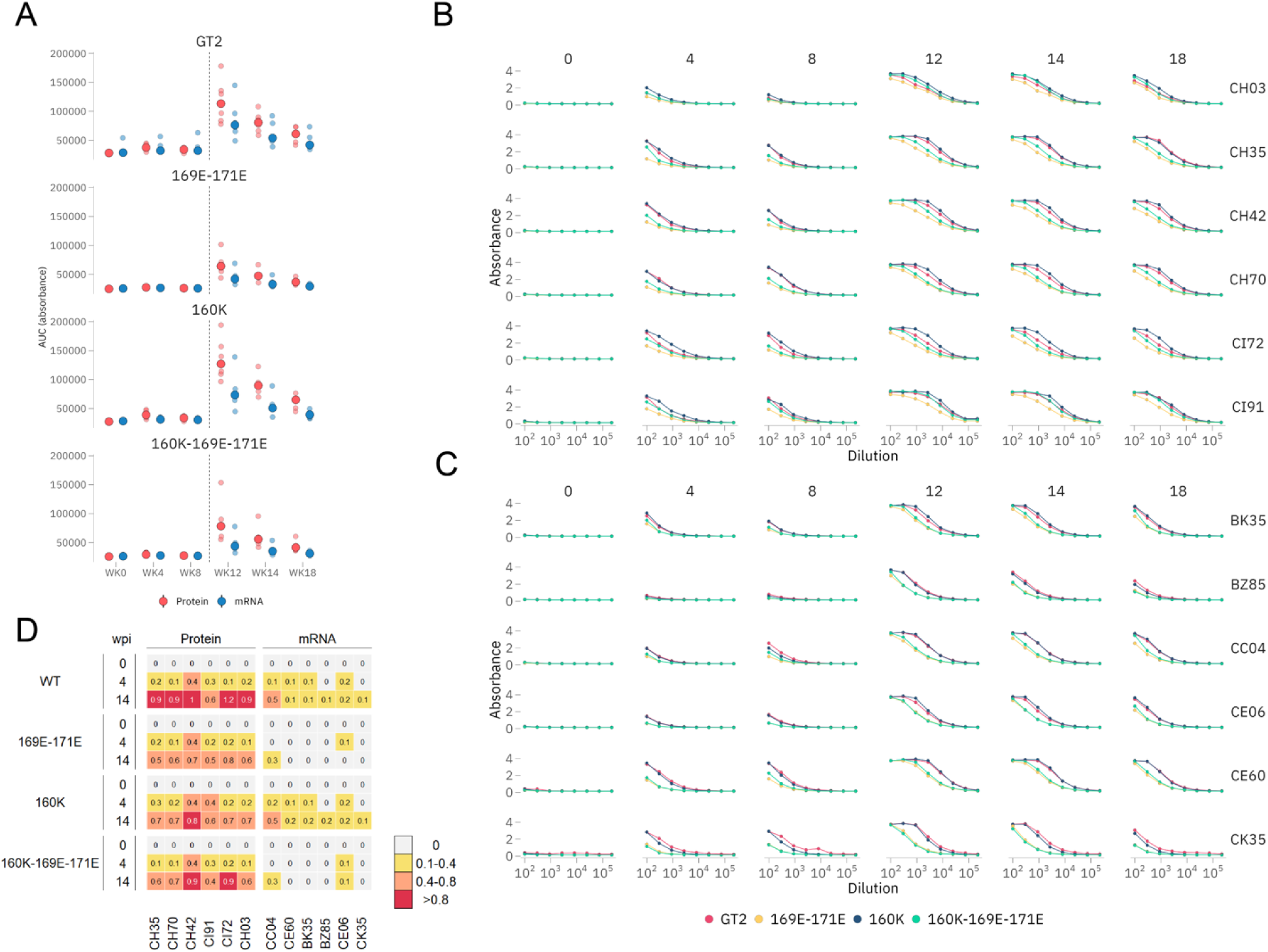
Longitudinal analysis of antigen and V2-apex epitope specific serum antibody responses in Q23-APEX-GT2 protein and mRNA vaccinated rhesus monkeys. Related to Figure 3. A. Binding of sera from immunized rhesus macaques to Q23-APEX-GT2 was evaluated using Enzyme-Linked Immunosorbent Assay (ELISA) starting at a serum dilution of 1:100. The data shows the serum titers over time, with a substantial increase in binding titers observed after the boost at week 10, indicated by the dotted line (panel A) with associated curves shown in panels B and C. Both the protein and mRNA immunized groups demonstrated increased titers post-boost. However, the protein-immunized group exhibited superior binding titers compared to the mRNA group, and both groups showed a decrease in titers as the study progressed, suggesting a waning immune response over time. B. Binding against Q23-APEX-GT2 and its epitope knockout variants (169E-171E, 160K, 160K-169E, 169E) from protein immunized rhesus macaques were tested with ELISA starting at serum dilution of 1:100 at week 0 (pre-immunization), 4, 8, 12, 14, 18. Absorbance at OD450 is shown. C. Binding against Q23-APEX-GT2 and its epitope knockout variants (169E-171E, 160K, 160K-169E, 169E) from mRNA immunized rhesus macaques were tested with ELISA starting at a serum dilution of 1:100 at week 0 (pre-immunization), 4, 8, 12, 14, 18. Absorbance at OD450 is shown. D. Biolayer Interferometry (BLI) was used to assess serum polyclonal IgG binding to Q23-APEX-GT2 and its epitope knockout variants (169E-171E, 160K, 160K-169E-171E) at weeks 0, 4, and 14. Both immunization groups developed Q23-APEX-GT2-specific antibody responses, with titers significantly higher in the protein-immunized group compared to the mRNA-immunized group. Epitope-dependent IgG responses were observed in both groups, with binding kinetics showing greater sensitivity to epitope knockouts, particularly in the mRNA group. Notably, the mRNA-immunized group exhibited a stronger reliance on specific epitope recognition, likely due to its lower overall polyclonal antibody titers compared to the protein-immunized group.

**Figure S6.**
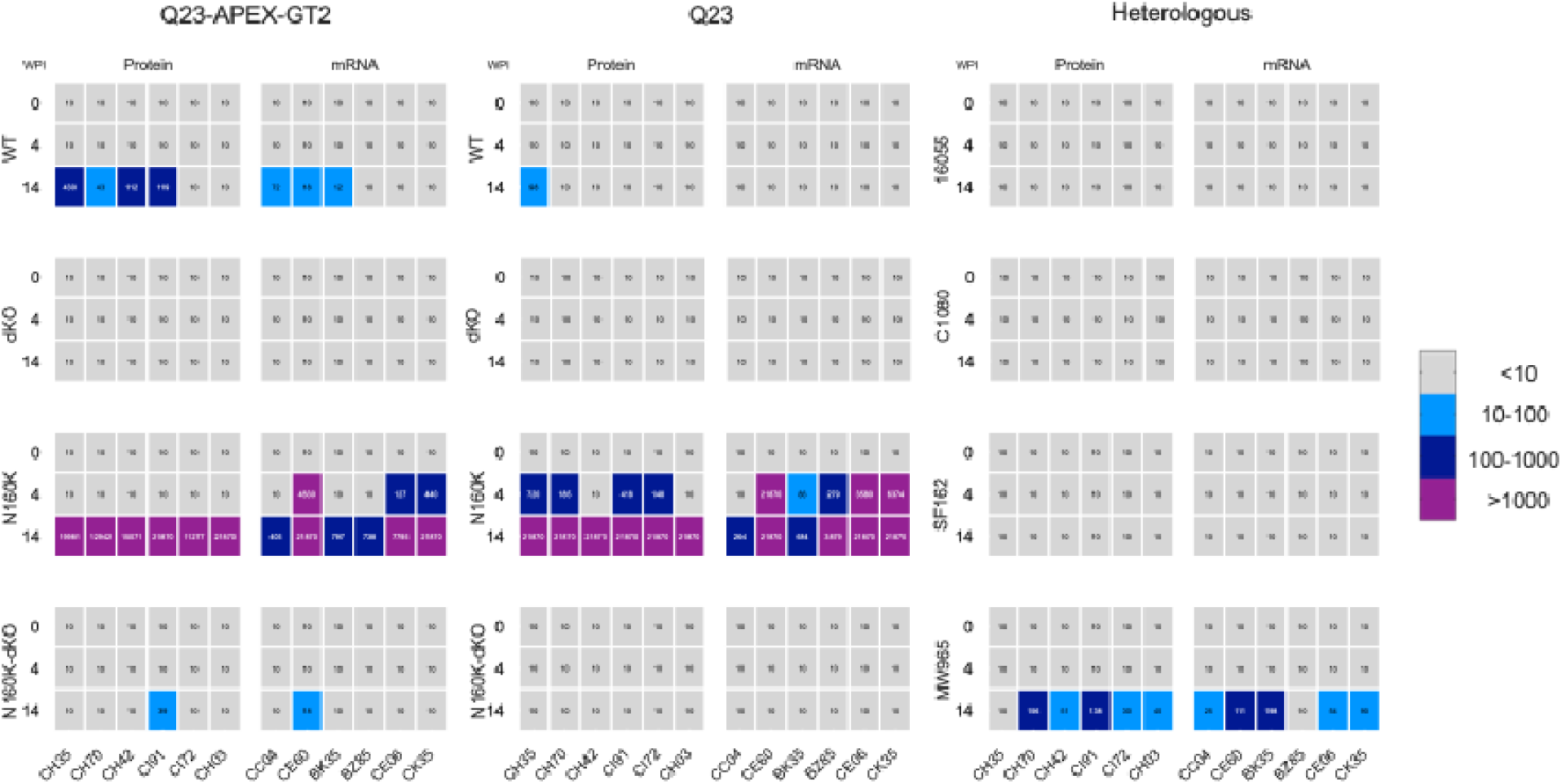
Neutralization by Q23-APEX-GT2 elicited rhesus immune serum antibody responses. Related to Figure 3. Heatmap depicting serum neutralization of pseudoviruses carrying Q23-APEX-GT2 mutations (132R-158T), the Q23.17 WT virus, and various V2-apex bnAb epitope variants (169E-171E [dKO], 160K, and 160K-169E-171E [tKO]) using immune sera collected at weeks 0, 4, and 14 from Q23-APEX-GT2 protein- or mRNA-LNP-vaccinated monkeys. Neutralization was dependent on strand C residues, as evidenced by the loss of neutralization with the 169E-171E variant. Notably, immune sera exhibited increased neutralization upon 160K glycan knockout, which was completely abrogated when strand C epitopes were disrupted in the N160K backbone (160K-169E-171E), reinforcing the role of strand C in V2-apex-targeted neutralization. Neutralization was also assessed against heterologous tier 2 HIV isolates (16055, C1080, and chimpanzee CAM13RRK, which shares the V2-apex site with HIV). Except for the protein-vaccinated monkey CH35, which exhibited some neutralization against CAM13KKR virus, no neutralization was detected. Additionally, two tier 1 viruses, SF162 and MW965, were tested. While most immune sera from both groups neutralized MW965, no neutralization was observed against SF162.

**Figure S7.**
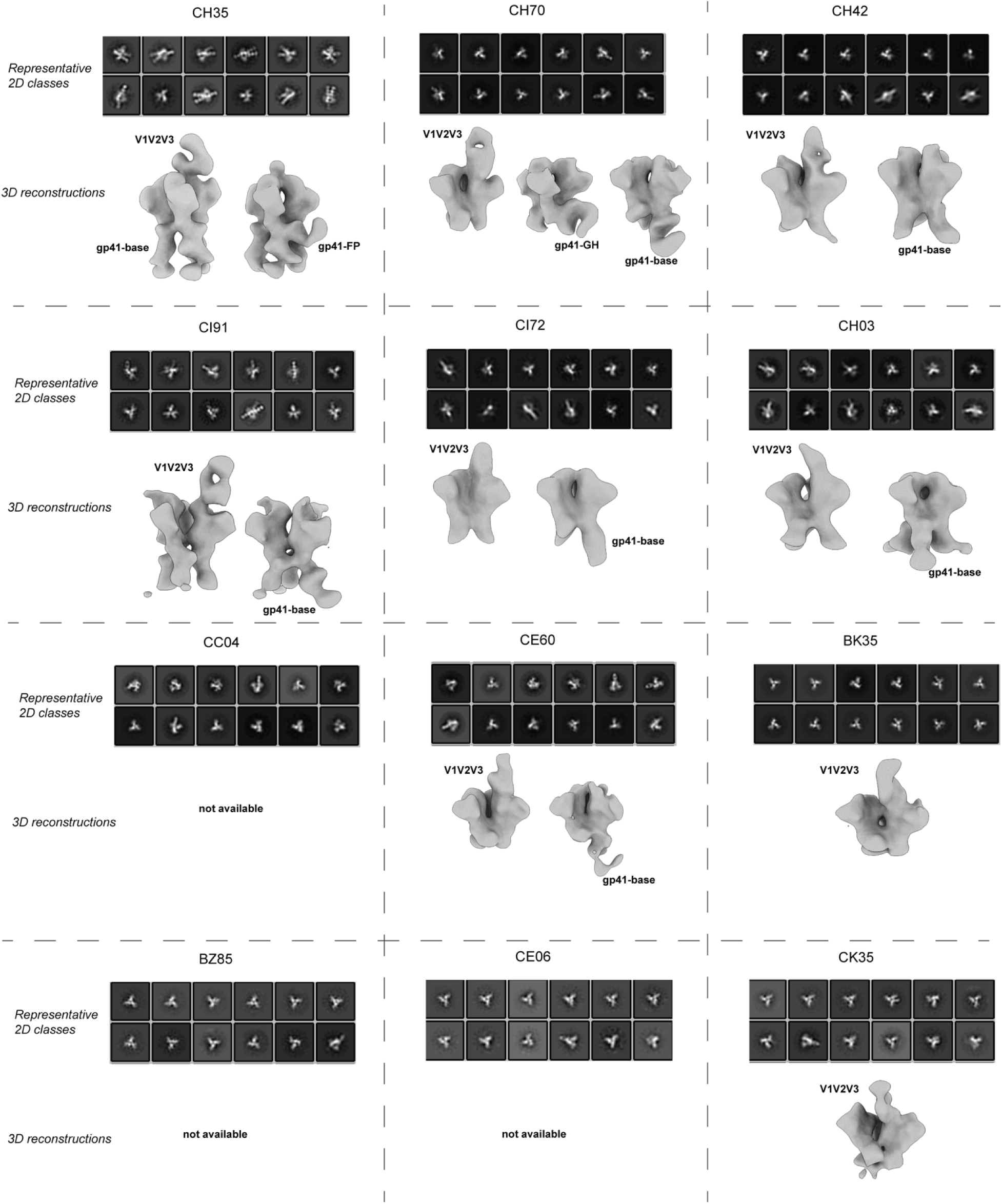
EMPEM 2D classes and 3D reconstructions. Related to Figure 3. Representative 2D classes of all immune complexes and final 3D reconstructions used to generate the segmented composite maps in Figure 3E are displayed for all 12 EMPEM samples. For datasets where too few Fab-bound particles were present for accurate 3D reconstruction, the label “not available” is used.

**Figure S8.**
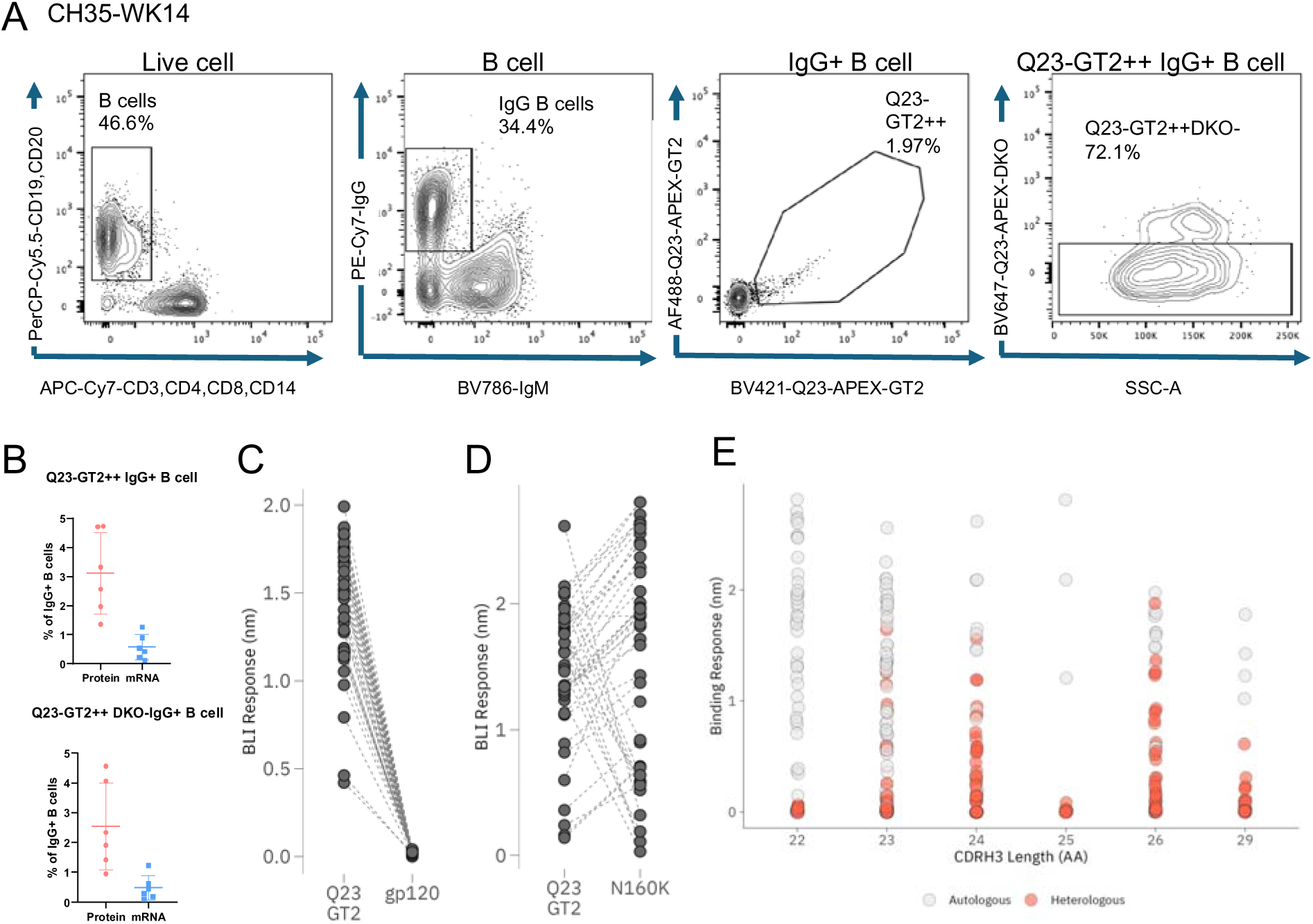
Antigen and epitope-specific IgG B cell profiling and mAb characterization in Q23-APEX-GT2 protein and mRNA-vaccinated monkeys. Related to Figures 4 and 5. A. Representative flow cytometry plots depicting Q23-APEX-GT2 antigen specific or V2-apex epitope specific (Q23-APEX-GT2 positive and Q23-APEX-GT2-dKO, R169E-K171E double knockout mutant) IgG B cells in week 14 lymph nodes of Q23-APEX-GT2 trimer protein-immunized CH35 rhesus macaque (RM). B. Scatter plots show the percentage of antigen-specific B cells (Q23-APEX-GT2⁺⁺) (upper panel) and epitope-specific B cells (Q23-APEX-GT2⁺⁺DKO⁻) (lower panel) among IgG⁺ B cells. The protein-immunized group is shown in red; and the mRNA-immunized group is shown in blue. C. BLI binding of isolated monoclonals from protein-immunized animals against the immunogen-matched protein trimer, Q23-APEX-GT2 (marked as GT2) and its monomeric gp120 protein. Complete loss of binding with gp120 monomers show trimer dependence of isolated antibodies. D. BLI binding of isolated monoclonals from protein-immunized animals against the immunizing protein Q23-APEX-GT2 (marked as GT2) and its N160K glycan knockout variant. Varying dependence on N160 glycan was seen with some mAbs showing reduced binding while several mAbs showed enhanced binding in the absence of N160 glycan. Several mAbs had little impact on binding in the absence of N160 glycan. E. BLI binding of isolated mAbs against autologous Q23-APEX-GT2 and heterologous HIV trimers from figure 5A. Each dot represents an antigen with heterologous binding only seen with mAbs of CDRH3 length 23AA and longer.

**Figure S9.**
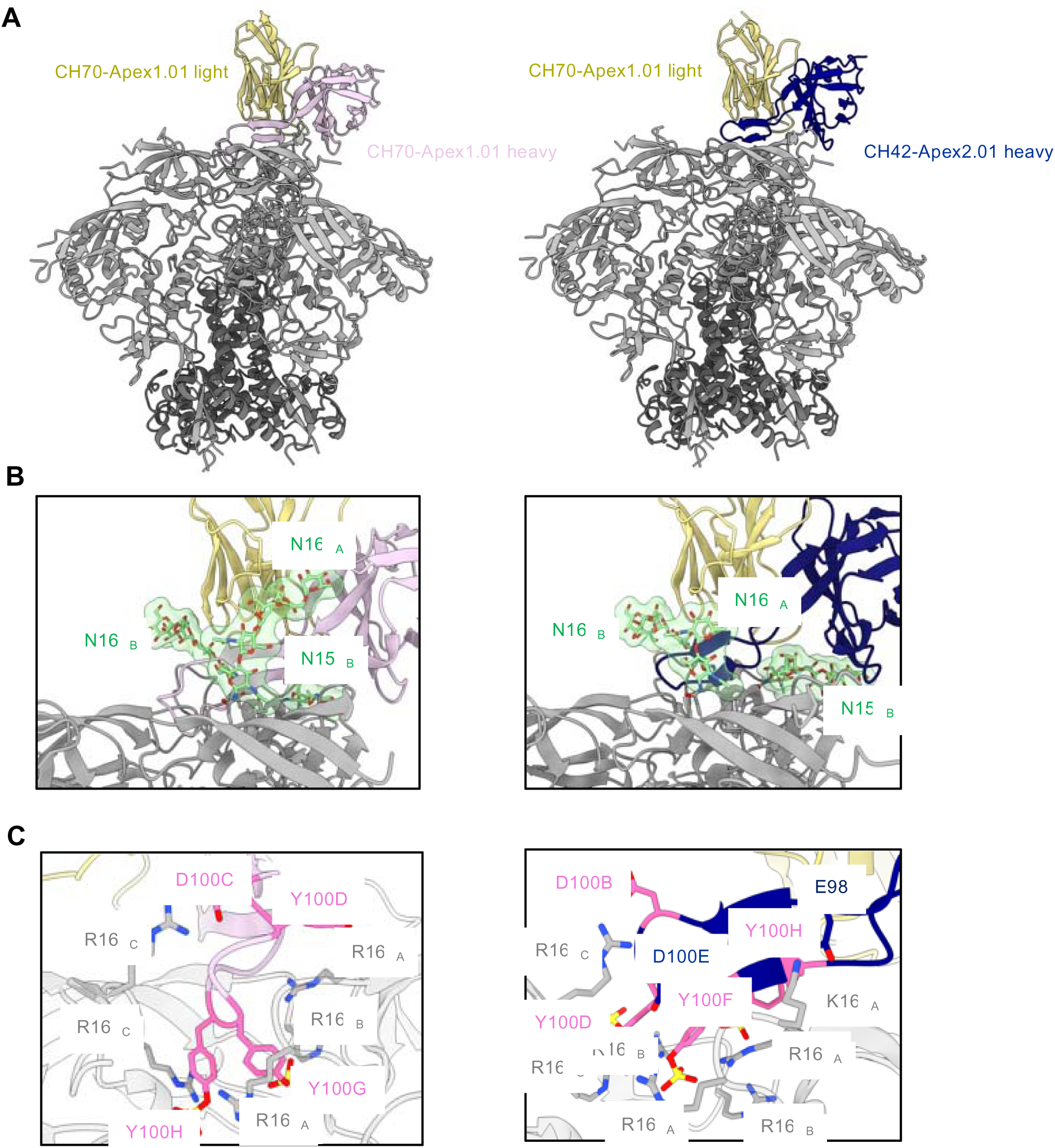
Cryo-EM structures of CH70-Apex1.01 and CH42-Apex2.01 in complex with Q23-APEX-GT2. Related to Figure 6. A. Overall structure of each respective lineage Fab in complex with the Q23-APEX-GT2 envelope trimer. B. Expanded interface view from panel (a) to highlight binding position and interactions with apical envelope glycans. Glycans bound by each respective Fab are shown in stick representation with transparent surfaces. C. Germline-encoded D3-15 residue interactions (dark pink) with core V2-apex epitope residues.

**Figure S10.**
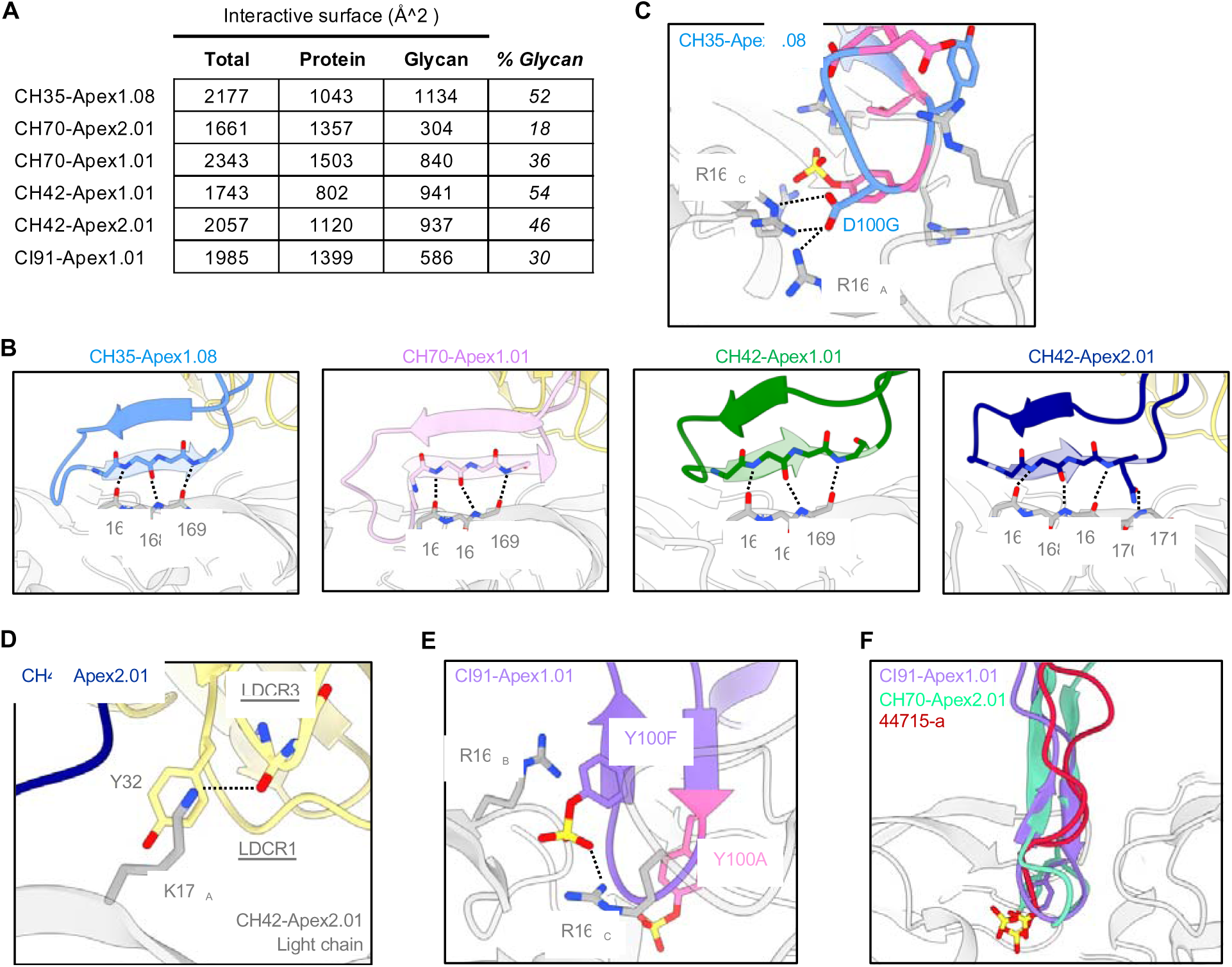
Select structural features of vaccine-elicited V2-apex targeted antibody lineages. Related to Figure 6. A. Quantification of the interactive surfaces for Fab:envelope cryo-EM structures determined in this study. B. Hydrogen bonding with the mainchain of V2 apex C-strand residues by combined-mode and axe-like lineages. C. CH35-Apex1.08 somatic mutant residue D100G (Y to D mutation) forms salt bridges with R166 and R169 from protomers A and C, respectively, completing recognition of these residue positions on all three protomers. D. CH42-Apex2.01 engages C-strand residue K171 through mainchain LCDR3 hydrogen-bonding and LCDR1 hydrophobic interactions mediated by Y32. E. CI91-Apex1.01 engages R166 and R169 from protomers C and A, respectively, via a second non-germline encoded sulfated tyrosine residue. The germline-encoded sulfated tyrosine residue, Y100A, is shown in pink for reference. F. Structural superimposition of needle-like vaccine-elicited lineages with SHIV-elicited mature bnAb lineage 44715-a (PDB-9bnm) by gp120 alignment reveals sulfated tyrosines at the HCDR3 tips each align within the middle of the trimer.

**Figure S11.**
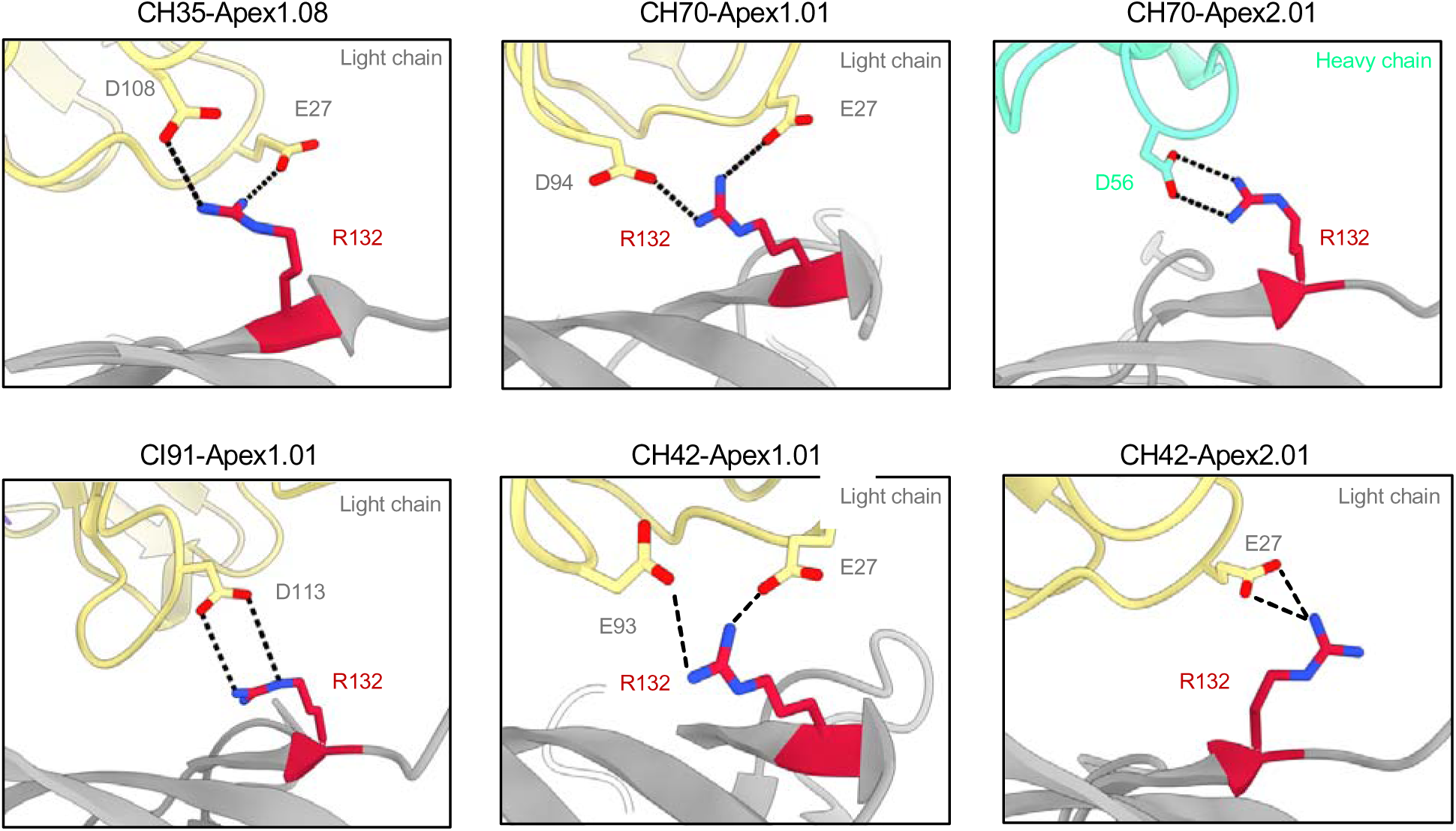
All vaccine-elicited lineages recognize the GT2 modification T132R. Related to Figure 6. Salt bridges formed between GT2 modification T132R and anionic residues from each vaccine-elicited lineage. Except for CH70-Apex2.01, all other lineages recognize T132R via the light chain.

**Figure S12.**
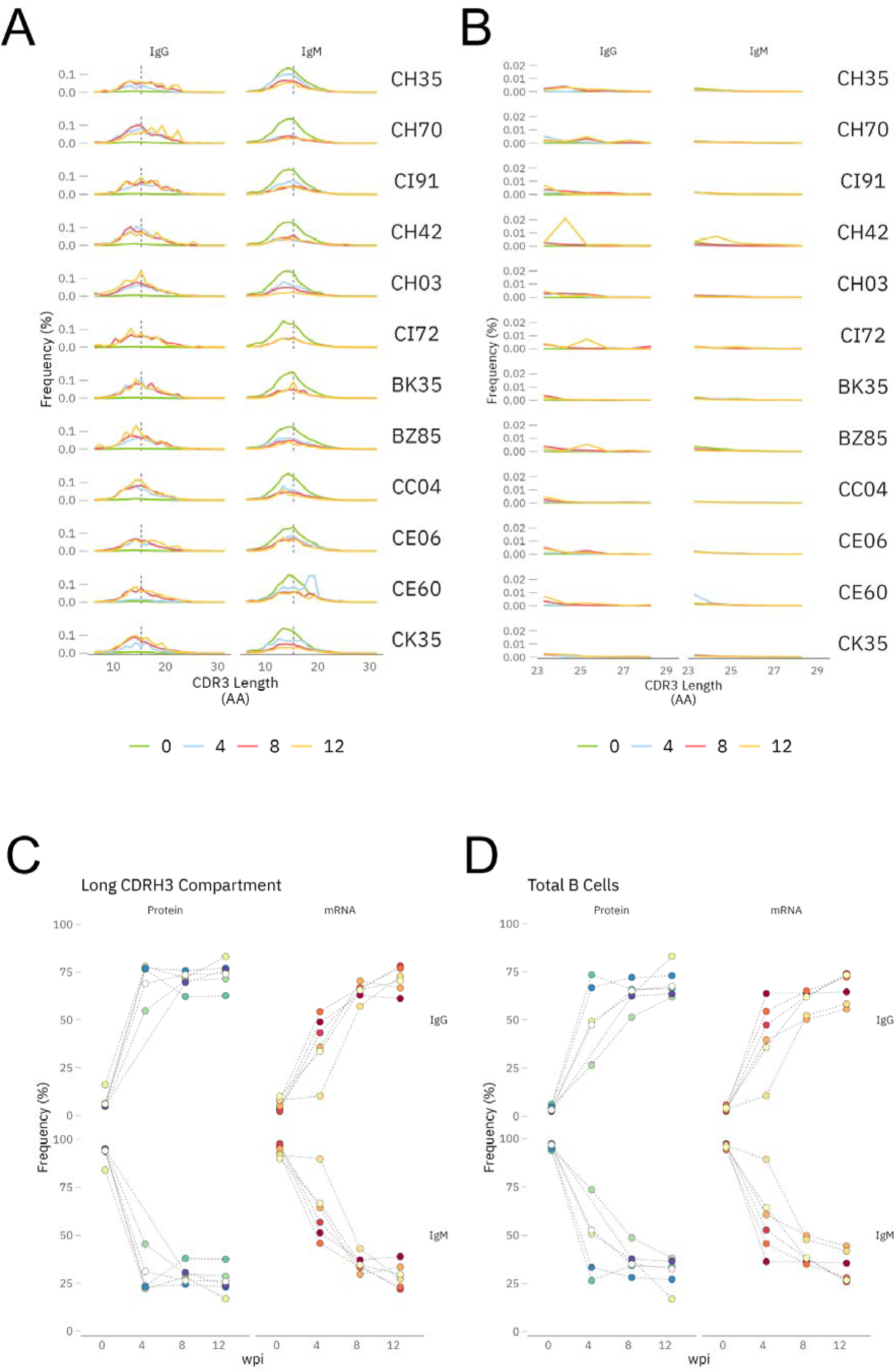
Longitudinal bulk BCR repertoire analysis of Q23-APEX-GT2 trimer immunized macaques show substantial expansion of long CDRH3 (≥23 amino acids) IgG B cell responses. Related to Figure 7. A. Frequency distribution of CDRH3 lengths in the bulk BCR repertoire of each rhesus macaque at pre-immunization (week 0) and post-immunization timepoints (weeks 4, 8, and 12). Data are shown separately for the IgG and IgM compartments. The distribution at week 0 represents the baseline repertoire before antigen exposure, while post-immunization timepoints reflect the evolution of B cell responses in responses to immunization. B. A magnified view of the subset of long CDRH3 sequences ≥23 amino acids (AA) from panel A, highlighting their enrichment within the IgG compartment across all animals. Compared to the pre-immunization repertoire, longer CDRH3 sequences show an increased frequency in the IgG compartment over time, suggesting preferential selection and expansion of class-switched IgG B cell clones with long CDRH3 loops following immunization. C. The frequency of class-switched B-cell receptors (BCRs) within the long CDRH3 compartment (≥23 AA) was analyzed in all immunized rhesus macaques, comparing protein and mRNA immunization regimens. Protein-immunized animals exhibited a substantial early shift towards IgG+ BCRs within the long CDRH3 subset by week 4, with levels stabilizing through weeks 8 and 12. In contrast, mRNA-immunized animals displayed a more gradual increase in IgG+ BCRs within the long CDRH3 compartment, continuing to rise steadily until week 12, suggesting a delayed but progressive class-switching response in this group. D. The frequency of class-switched BCRs in the total B-cell compartment demonstrated more heterogeneous profiles of IgG and IgM across individual animals, irrespective of immunization type. This variability highlights differential class-switching kinetics at the total B-cell level compared to the long CDRH3-specific compartment.

**Figure S13.**
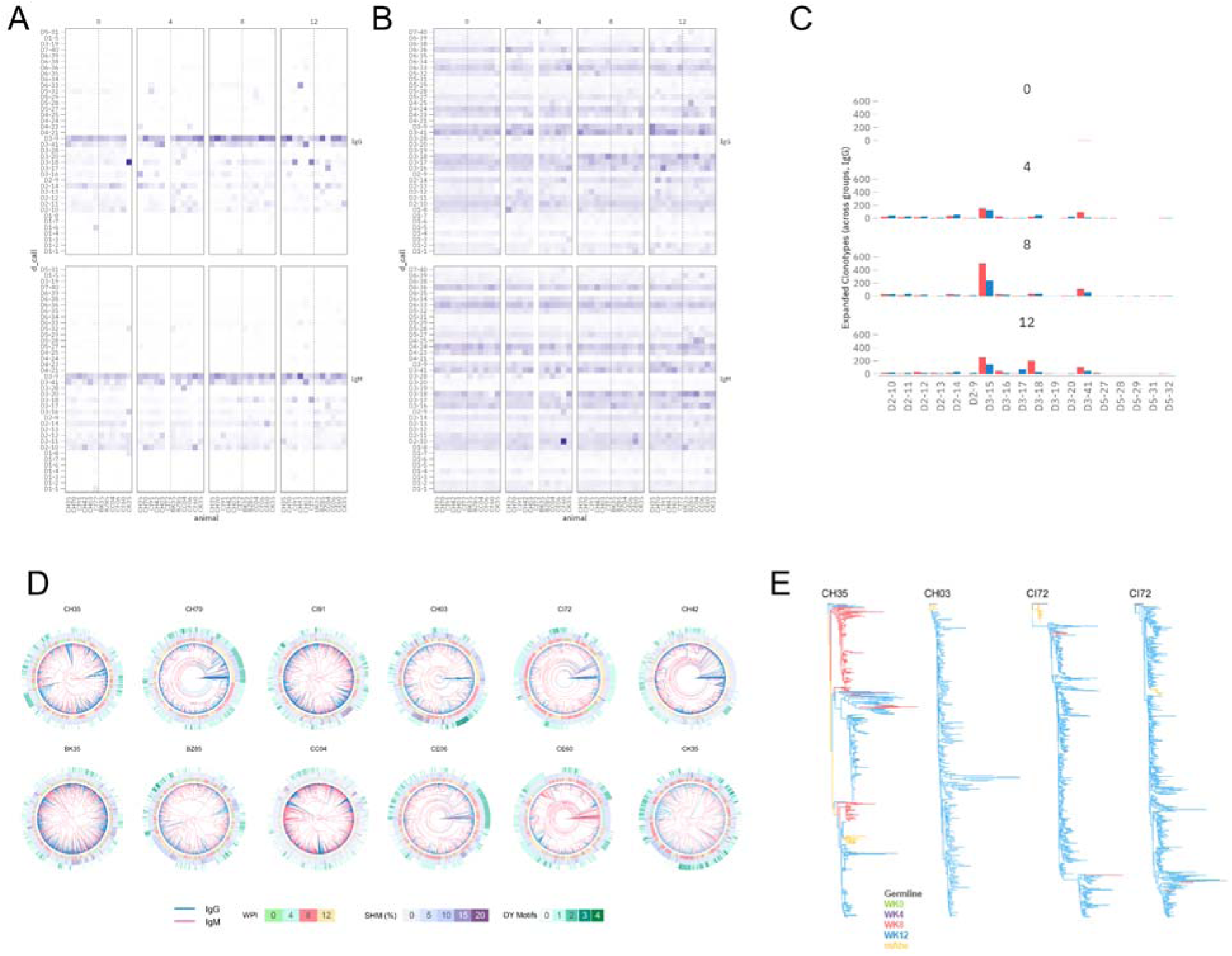
Longitudinal analysis of D-gene distribution, IgG class switching, and lineage expansions in long CDRH3 B cell responses in Q23-APEX-GT2 trimer immunized macaques. Related to Figure 7. A – B. Frequency distribution of D-gene usage within B cell receptors (BCRs) categorized by long (≥23 AA) or ≤22 AA complementarity-determining region 3 of the heavy chain (CDRH3). A substantial enrichment of the IGHD3 gene family is observed across all animals, suggesting a preferential selection or expansion of this gene family in response to immunization. The distribution pattern indicates a bias toward specific D-gene segments contributing to the formation of longer or shorter CDRH3 loops, potentially influencing antigen binding characteristics. C. D-gene usage within expanded clonotypes in Q23-APEX-GT2 protein (red) and mRNA (blue) group immunized animals highlights a notable increase in the frequency of IGHD3-15, IGHD3-18, and IGHD3-41 across both immunization groups. The differential usage of additional D-genes suggests a level of variability in BCR repertoire formation, potentially driven by antigen engagement and selection pressures unique to each immunization strategy. These findings imply that while certain D-genes are commonly enriched, the breadth of D-gene utilization varies between protein- and mRNA-based immunization. D. Circular phylogenetic trees displaying expanded clonotypes with associated genotypic data for all immunized rhesus macaques. Each tree represents the evolutionary relationships among B cell receptor (BCR) sequences within individual animals over the immunization period. Branch lengths correspond to sequence divergence, with nodes colored according to immunoglobulin isotype (IgM, IgG). In protein-immunized animals, we observe a clear longitudinal shift towards IgG-dominated clonotypes, indicative of class-switch recombination and affinity maturation. This transition is accompanied by increasing somatic hypermutation (SHM) levels and a progressive enrichment of the anionic sulfation motif (DY) within the complementarity-determining regions (CDRs), suggesting selective pressure favoring V2-apex epitope engagement and maturation. In contrast, mRNA-immunized animals exhibit a distinct pattern, with expanded clonotypes remaining predominantly IgM-expressing, and fewer signs of advanced maturation. The limited increase in SHM and minimal enrichment of sulfation motifs imply a reduced capacity for V2-apex directed clonotype evolution. E. Phylogenetic trees depicting additional monoclonal antibodies (mAbs) from Figure 4F illustrate the presence of lineage members as early as week 8, indicating that these mAbs were seeded by the Q23-APEX-GT2 prime. The branching patterns and sequence relationships confirm the early establishment and diversification of the lineage following priming.

**Figure S14.**
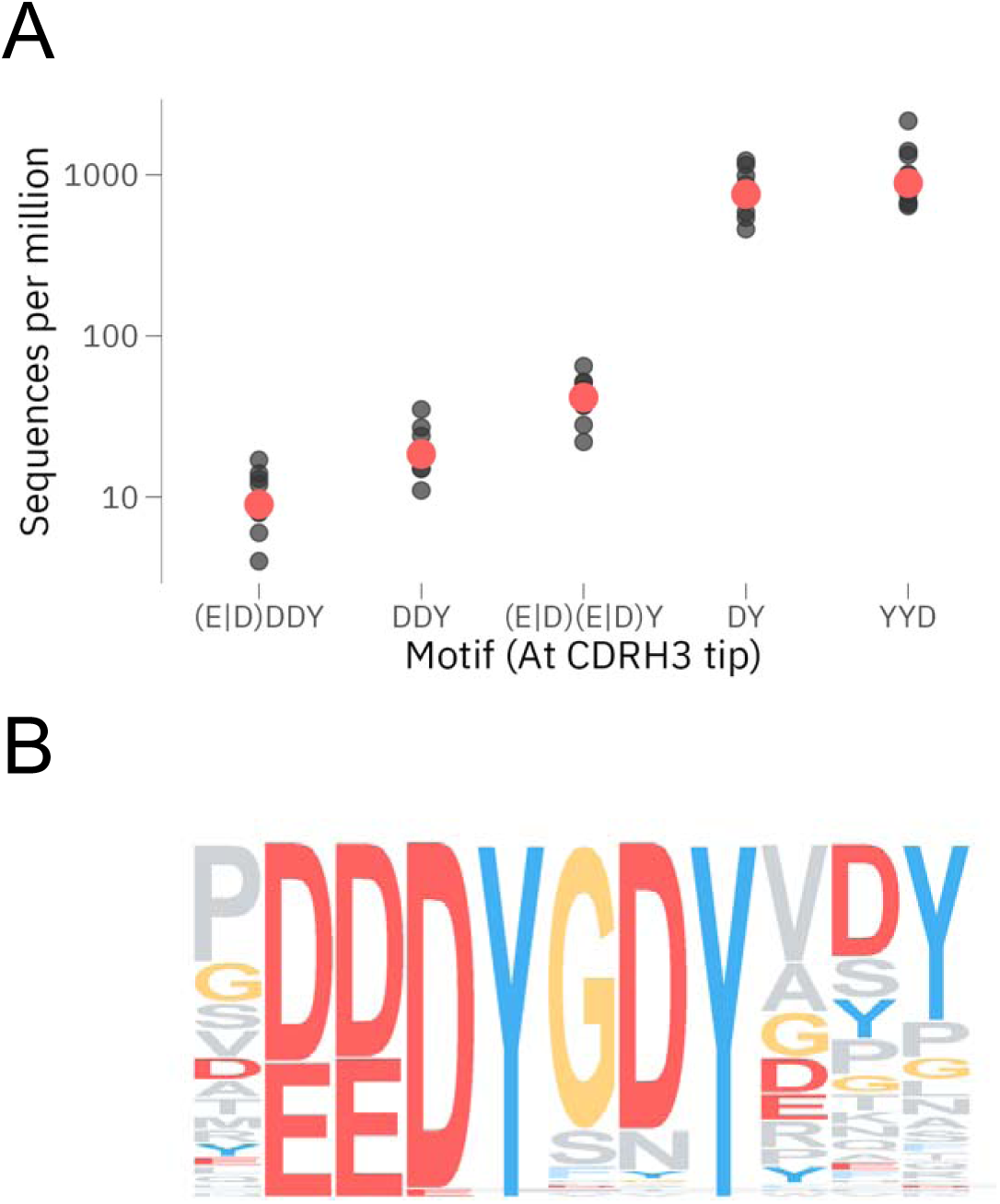
Frequency of long CDRH3 V2-apex bnAb-like precursors in naïve human B cell repertoires and commonalities with rhesus bnAb precursors. Related to Figure 7. A. The frequency of charged motifs at the apical position in long CDRH3 loops (≥23 amino acids) within naïve human BCR repertoires is shown as sequences per million. The data demonstrates an increasing occurrence of these motifs as the stringency of the charged residues decreases. Specifically, the DY and YYD motifs appear more frequently than motifs containing an additional charged residue, such as EDDY or DDDY, suggesting selective constraints on excessive charge at the CDRH3 apex. B. A sequence logo representation of human naïve BCR sequences containing motifs similar to the rhesus macaque IGHD3-15 germline-derived sequence (YEDDYGYYT). The sequence logo highlights the frequent presence of the EDDYG motif, while the downstream YYT residues appear less commonly, indicating that the YYT extension may not be as strongly conserved within the human repertoire. This suggests commonalities and potential differences that may influence function and antigen driven selection between human and rhesus BCR repertoires.

